# Transcriptome analysis of apple leaves infected by the rust fungus *Gymnosporangium yamadae* at two sporulation stages (spermogonia and aecia) reveals specific host responses, rust pathogenesis-related genes and a shift in the phyllosphere fungal community composition

**DOI:** 10.1101/717058

**Authors:** Si-Qi Tao, Lucas Auer, Emmanuelle Morin, Ying-Mei Liang, Sébastien Duplessis

## Abstract

Apple rust disease caused by *Gymnosporangium yamadae* is one of the major threats to apple orchards. In this study, dual RNA-seq analysis was conducted to simultaneously monitor gene expression profiles of *G. yamadae* and infected apple leaves during the formation of rust spermogonia and aecia. The molecular mechanisms underlying this compatible interaction at 10 and 30 days post inoculation (dpi) indicate a significant reaction from the host plant and comprise detoxication pathways at the earliest stage and the induction of secondary metabolism related pathways at 30dpi. Such host reactions have been previously reported in other rust pathosystems and may represent a general reaction to rust infection. *G. yamadae* transcript profiling indicates a conserved genetic program in spermogonia and aecia that is shared with other rust fungi, whereas secretome prediction reveals the presence of specific secreted candidate effector proteins expressed during apple infection. Unexpectedly, the survey of fungal unigenes in the transcriptome assemblies of inoculated and mock-inoculated apple leaves reveals that *G. yamadae* infection modifies the fungal community composition in the apple phyllosphere at 30 dpi. Collectively, our results provide novel insights into the compatible apple-apple rust interaction and advance the knowledge of this heteroecious demicyclic rust fungus.

## Introduction

Apple (*Malus domestica* Borkh.), one of the major fruitful and economic fruiters in temperate regions of the world, is susceptible to the rust pathogen *Gymnosporangium yamadae* (Cummins and Hiratsuka 2003; Kern 1973; Peterson 1967). Heavy infection can lead to significant decreases in fruit yield and economic losses in most apple-planting regions in Asia (Kim and Kim 1980; Wang et al. 2010). As a heteroecious and demicyclic rust fungus, *G. yamadae* produces four morphologically different spores on two taxonomically different hosts (*Malus* spp. and *Juniperus chinensis*) to complete its life cycle (Kern et al. 1973; Yun et al. 2005). In early spring, telia germinate and produce gelatinous tendrils containing haploid basidiospores which can disperse into the air. The released spores parasitized the surface of *Malus* leaves and successful infection leads to orange-yellow spots on the upper surface. Then the spots turn to bright orange-red with a red border and may exhibit small raised black dots in the centre of the spots. These lesions grow through the leaf and develop small brownish and spiky projections on the lower surface of leaf (Cummins and Hiratsuka 2003).

In Asia, junipers are widely cultivated and frequently found in graveyards, parks and roadsides. Trans-provincial transportation of juniper seedlings with overwintering fungal infections increases the risk of disease occurrence (Tao et al. 2018). Twenty *Malus* spp. have been tested for disease resistance and only *M. halliana* showed high resistance to *G. yamadae* basidiospores, the others all are highly susceptible (Harada 1984). To control this disease, protective fungicides are applied on apple leaves to prevent penetration of basidiospores into the host (Guo 1994), and the control efficiency is influenced by the spraying timing. Still, no effective management strategy has been established because the molecular mechanisms underlying the apple-apple rust interaction has not been investigated, and the fungal parasitic factors still remain unknown.

In the past few years, transcriptomic and bioinformatic approaches have allowed the identification of candidate virulence factors of many rust fungi at different infection or life-cycle stages (Lorrain et al. 2019). Similarly, efforts have been made for the identification of host defense mechanisms activated in response to rust infection at a transcriptomic scale (Rinaldi et al. 2007; Schneider et al. 2011; Ullah et al. 2019; van de Mortel et al. 2007). Dual RNA-seq has been applied to simultaneously detect gene expression changes in both host and pathogens, including plant-rust fungi interactions (Dobon et al. 2016; Fernandez et al. 2012; Kawahara et al. 2012; Teixeira et al. 2014; Westermann et al. 2017). Such studies provided invaluable resources to help identifying host molecular alterations during direct fungal action and better understand the strategies used by the pathogen to manipulate the host during the infection process.

In addition to pathogenic fungi, the plant phyllosphere harbours large numbers of other microbiota, such as bacteria, non-pathogenic fungi and archaea (Lindow et al. 2003; Vorholt 2012). Among these microbiotas, some play a direct role in protecting their host against pathogens (Hassani et al. 2018). Massive parallel sequencing technologies are remarkable tools to describe the in-depth composition and structure of microbial communities associated with leaves of many plants, including apple (Agler et al. 2016; Becker et al. 2008; Camatti-Sartori et al. 2005; Leveau and Tech 2011; Reisberg et al. 2012). However, the systematic exploration of phyllosphere fungal communities remains limited and only a few reports have shown the microbes from the plant phyllosphere could play an important role in resistance to pathogens (Busby et al. 2016; Hassani et al. 2018; Ritpitakphong et al. 2016; Vogel et al. 2016). The study of microbiome in the plant phyllosphere during rust infection is barely explored (Busby et al. 2016).

In this study, we conducted a dual RNA-seq analysis of apple leaves inoculated with the rust fungus *G. yamadae* at two time points, 10 days post inoculation (10 dpi) and 30 days post inoculation (30 dpi), compared to mock-inoculated treatment. We aimed to explore the host responses to *G. yamadae* infection and to identify rust genes expressed during the interaction with apple leaves, with an emphasis on secreted proteins which may relate to pathogenicity. We also report on the impact of *G. yamadae* infection on fungal communities of the apple phyllosphere.

## Results

### Experimental design and RNA-seq results

Apple rust spermogonia and aecia were collected from leaves of two-year-old apple seedlings inoculated with *G. yamadae* basidiospores after 10 days post-inoculation (dpi) and 30 dpi, respectively (Figure 1). Control mock-inoculated treatments with sterile water were obtained from plants grown in strictly similar conditions at 10 and 30 dpi (Supplementary file: Figure S1). Three biological replicates were collected for each sample. The collected inoculated samples included the fungal sporulation structures and the leaf area discoloured during the fungal infection as visible on Figure 1. Leaf samples of similar area were collected for the mock-inoculated controls. To ensure the production of the two fungal sporulation structures targeted in this study, the infection procedure requires to keep the two-year-old apple seedlings bagged for 10 days in controlled green-house conditions before to move them outdoors until 30 days and aecia differentiation. This experimental set-up allows for direct comparisons between inoculated and mock-inoculated samples at the two stages, but environmental effects do not allow for direct comparison between time points.

**Figure 1.**
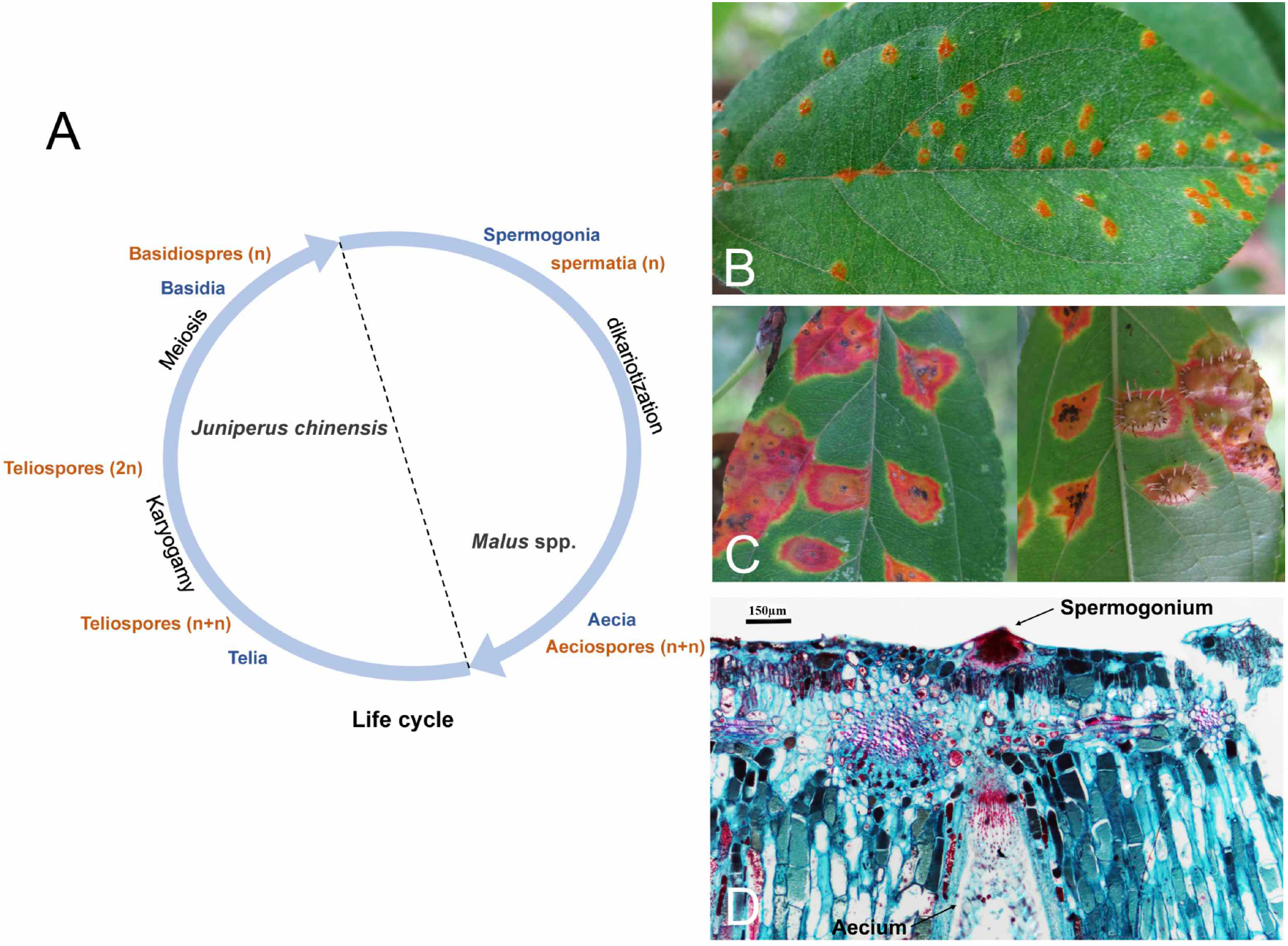
Spermogonia and aecia stages in the *Gymnosporangium yamadae* life cycle. A: schematic view of *G. yamadae* life cycle illustrating the spore stages used for inoculation (here teliospores and basidiospores from *Juniperus chinensis* as primary inoculum) and for samples collection (spermogonia and aecia on *Malus domestica*). B: Spermogonia visible on the upper surface of apple leaves 10 days after controlled inoculation. C: upper (left) and lower (right) sides of the same apple leaf are shown with coloured area around the sporulation zones and tubular aecia extruding on the lower siode of the leaf 30 days after inoculation. D: longitudinal section of a *G. yamadae* infected apple leaf stained with aniline blue and phenol red, showing a tubular aecium in red formed on the lower leaf epidermis, directly below a globoid spermogonium also in red, shown on the upper epidermis.

Total RNA for each replicate was isolated and sequenced using Illumina Hiseq platform and subjected to a dedicated analysis pipeline (Supplementary file: Figure S2). After removing reads of low quality, 264 and 269 million paired clean reads were obtained for infected apples leaves, and 194 and 151 million clean reads from healthy leaves, at 10dpi and 30dpi, respectively (Table 1). Paired-end reads were aligned to the *M. domestica* reference genome (Velasco et al. 2010) using Tophat v.2.0.12 (Trapnell et al. 2009) and the mapped rates are relatively lower in infected leaves than in healthy leaves and even lower at 30dpi (24.68% for average), which reflects the development of the rust pathogen inside the infected leaves (Table 1). Pearson correlation coefficients (*r*^2^) were calculated between biological replicates and conditions to assess the overall reproducibility of the data (Figure 2A). The results showed a strong correlation between replicates of a single condition, and a clear separation between independent conditions (Figure 2A). The principal components analysis based on read counts confirmed the distinction of expression profiles of inoculated and mock inoculated samples at the two time points and the proximity of biological replicates (Figure 2B).

**Table 1.** Sequencing and mapping information of RNA-seq data from 12 libraries of apple leaves.

| Sample | Library | Clean reads | Reads mapped to *Malus domestica* | Rust unigenes<sup>a</sup> |
| --- | --- | --- | --- | --- |
| Inoculated (10 dpi) | Inoculated (10 dpi)_1 | 100,206,930 | 61,059,189 (60.9%) | 30,293 |
|  | Inoculated (10 dpi)_2 | 77,191,936 | 49,422,953 (64.0%) |  |
|  | Inoculated (10 dpi)_3 | 87,378,970 | 56,459,826 (64.6%) |  |
|  | total | 264,777,836 | 166,941,968 (63.2%) |  |
| Mock-inoculated (10 dpi) | Mock-inoculated (10 dpi)_1 | 65,394,614 | 51,116,139 (78.2%) | - |
|  | Mock-inoculated (10 dpi)_2 | 66,237,512 | 51,749,324 (78.1%) |  |
|  | Mock-inoculated (10 dpi)_3 | 62,536,254 | 49,104,270 (78.5%) |  |
|  | total | 194,186,380 | 151,969,733 (78.3%) |  |
| Inoculated (30 dpi) | Inoculated (30 dpi)_1 | 87,373,062 | 22,244,421 (25.5%) | 22,717 |
|  | Inoculated (30 dpi)_2 | 97,680,330 | 19,448,833 (19.9%) |  |
|  | Inoculated (30 dpi)_3 | 84,796,366 | 24,319,732 (28.7%) |  |
|  | total | 269,849,758 | 66,012,986 (24.7%) |  |
| Mock-inoculated (30 dpi) | Mock-inoculated (30 dpi)_1 | 41,186,566 | 33,721,232 (81.9%) | - |
|  | Mock-inoculated (30 dpi)_2 | 51,906,804 | 42,616,916 (82.1%) |  |
|  | Mock-inoculated (30 dpi)_3 | 58,183,906 | 47,693,953 (81.9%) |  |
|  | total | 151,277,276 | 124,032,101 (82.0%) |  |

<sup>a</sup> Un-mapped reads from inoculated apple leaves at 10dpi and 30dpi were assembled and compared to Basidiomycota genomic data and Pucciniales ESTs to determine rust unigenes for spermogonia and aecia, respectively

**Figure 2.**
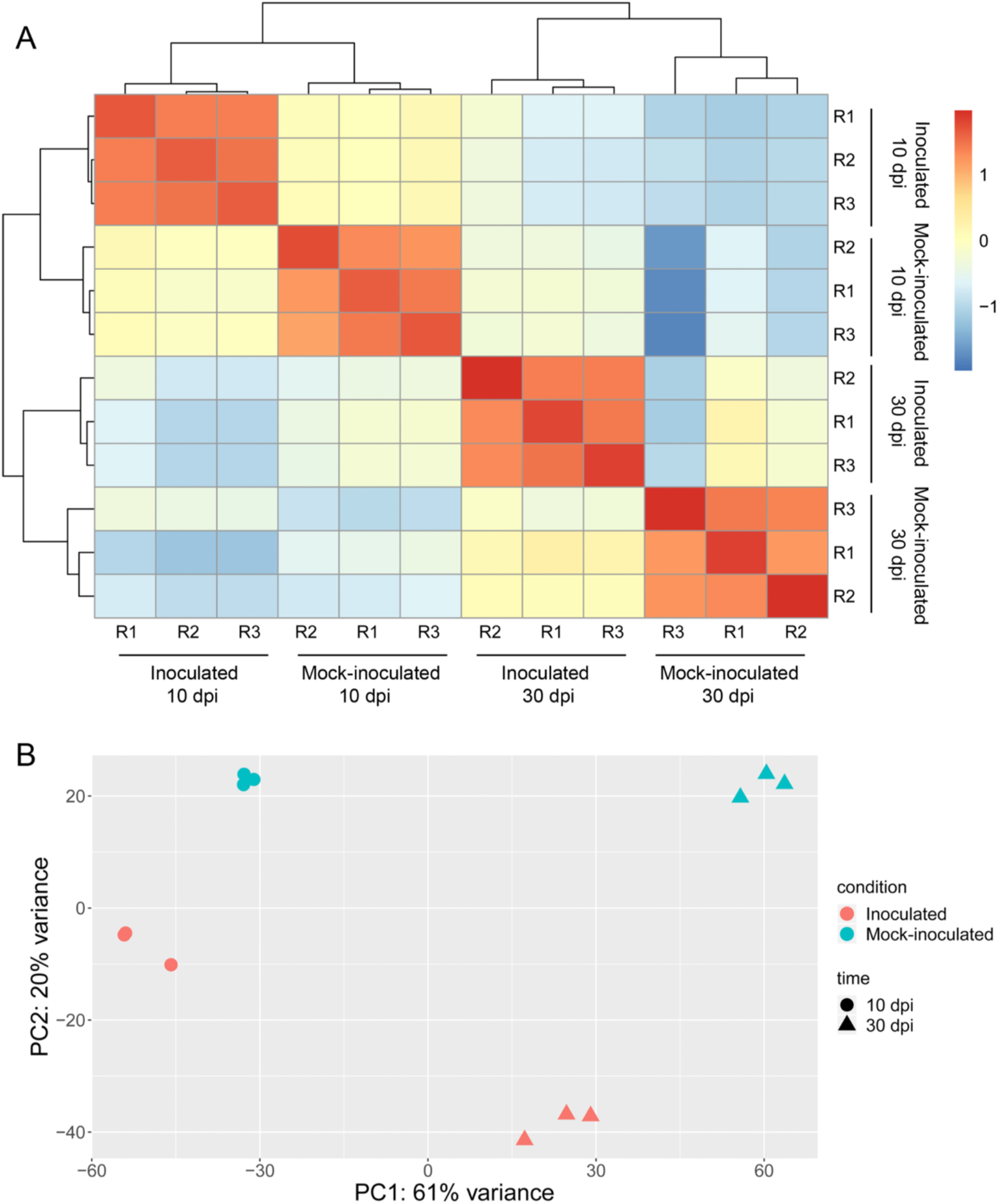
Assessment of RNA-seq data reproducibility. A: Hierarchical clustering of replicates of the inoculated and mock-inoculated conditions at the two time points, 10 and 30 days post-inoculation (dpi), based on Pearson correlation coefficients between samples. B: Principal component analysis (PCA) of read counts of inoculated and mock-inoculated conditions at the time points 10 and 30 dpi, showing the clear separation of the four tested conditions and the proximity of biological replicates. The two principal components explain 81% of the total variance.

After mapping the Illumina reads onto the 57,386 predicted genes in the apple genome (Velasco et al. 2010), 26,530 and 29,227 transcripts were identified from inoculated and mock inoculated conditions at 10 dpi, respectively. Besides, 29,725 and 26,867 transcripts were found in inoculated and mock inoculated conditions at 30 dpi, respectively (Supplementary file: Table S1). Reads unmapped to the apple genome in each inoculated condition were pooled and assembled using *de novo* assembly Trinity (Grabherr et al. 2011) and fragments per kilobase of transcript sequence per millions base pairs sequenced (FPKM) values were calculated for all assembled unigenes. In order to identify apple rust transcripts in absence of a reference genome for *G. yamadae*, these tentative fungal unigenes were compared to reference badisiomycete genomes, including four rust species and 97,034 Pucciniales ESTs. In total, 30,293 and 22,717 unigenes were considered as *G. yamadae* transcripts expressed in infected apple leaves at the spermogonial and aecial stage, respectively (Supplementary file: Figure S2, Table S2). After assignment of unigenes to the plant host and to the rust fungus, we considered unassigned reads and compared them to the NCBI database, revealing the presence of many fungal sequences belonging to the Ascomycota (data not shown). This observation prompted us to explore the fungal community composition in the apple phyllosphere in more details.

### Apple leaves transcriptional changes upon *G. yamadae* infection

The four transcriptomes obtained in this study were compared to investigate specific gene expression profiles in apple leaves during infection by *G. yamadae* at the spermogonial and aecial stages. A total of 34, 246 transcripts (59.7% of *M. domestica* genes) were expressed at least in one condition, and 22,167 (38.6%) transcripts were expressed in all conditions. We found 300 (0.9%) transcripts exclusive of the inoculated conditions and 256 (0.7%) of the mock inoculated conditions, whereas a total of 1,115 (3.3%) genes were only expressed at 10 dpi and 1,283 (3.7%) at 30 dpi (Figure 3). To explore the molecular mechanisms triggered in response to *G. yamadae* infection, pairwise comparisons of the host plant transcriptomes were conducted between inoculated and mock inoculated apple leaves at the two time points and transcripts showing significant levels of expression regulation (p_*adj*_ ≤ 0.05) were regarded as differentially expressed genes (DEGs). The comparisons between inoculated and mock inoculated apple leaves generated 13,961 DEGs at 10 dpi (7,784 up-regulated and 6,177 down-regulated) and 4,428 at 30 dpi (2,883 up-regulated and 1,545 down-regulated) (Supplementary file: Table S1, Figure 4). Hierarchical clustering of DEGs showed dynamic expression profiles at the two infection stages in this compatible apple-apple rust interaction (Figure 4). Among these DEGs, 579 and 291 were respectively up-regulated and down-regulated in response to the infection of *G. yamadae*, representing *G. yamadae* infection-related DEGs (Figure 4). Among highly induced apple transcripts in infected leaves at 10 dpi, 56 unigenes encoding glutathione S-transferase (GST) showed a fold-change up to 194, among which 23 had a fold-change over 10, whereas only 14 GSTs were up-regulated at 30 dpi with a fold-change ranging from 3 up to 27 (Supplementary file: Table S1). In total, 12 GST transcripts were induced both at 10 and 30 dpi. Interestingly, eight mannitol dehydrogenase-encoding genes were also among highly induced transcripts at 10 dpi (fold-change up to 231) while no significant change was observed at 30dpi. Several gene functions related to cell-wall, phytohormones, or response to pathogens were found among the top highly expressed DEGs at 10 and 30 dpi (Table 2).

**Table 2.**
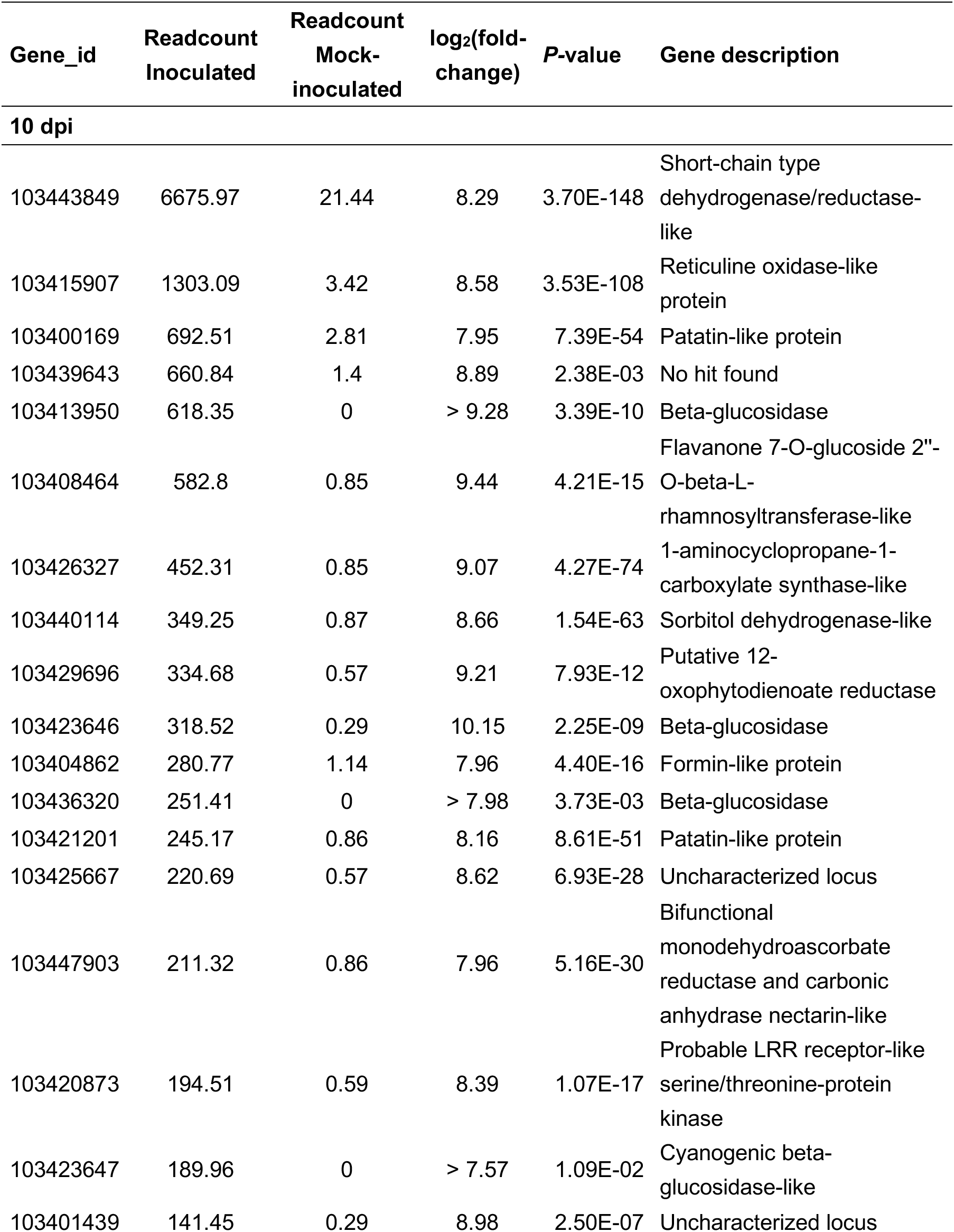
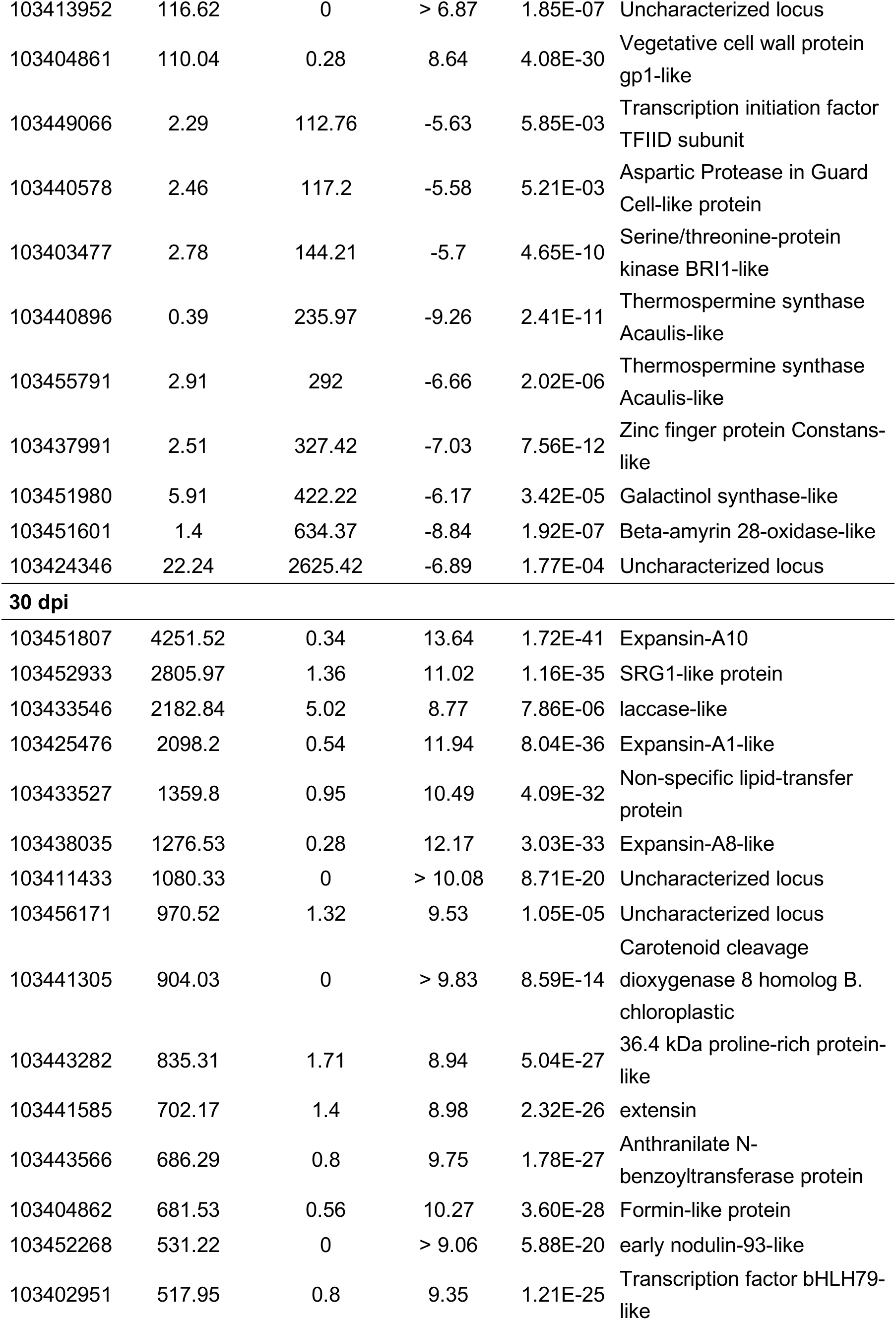
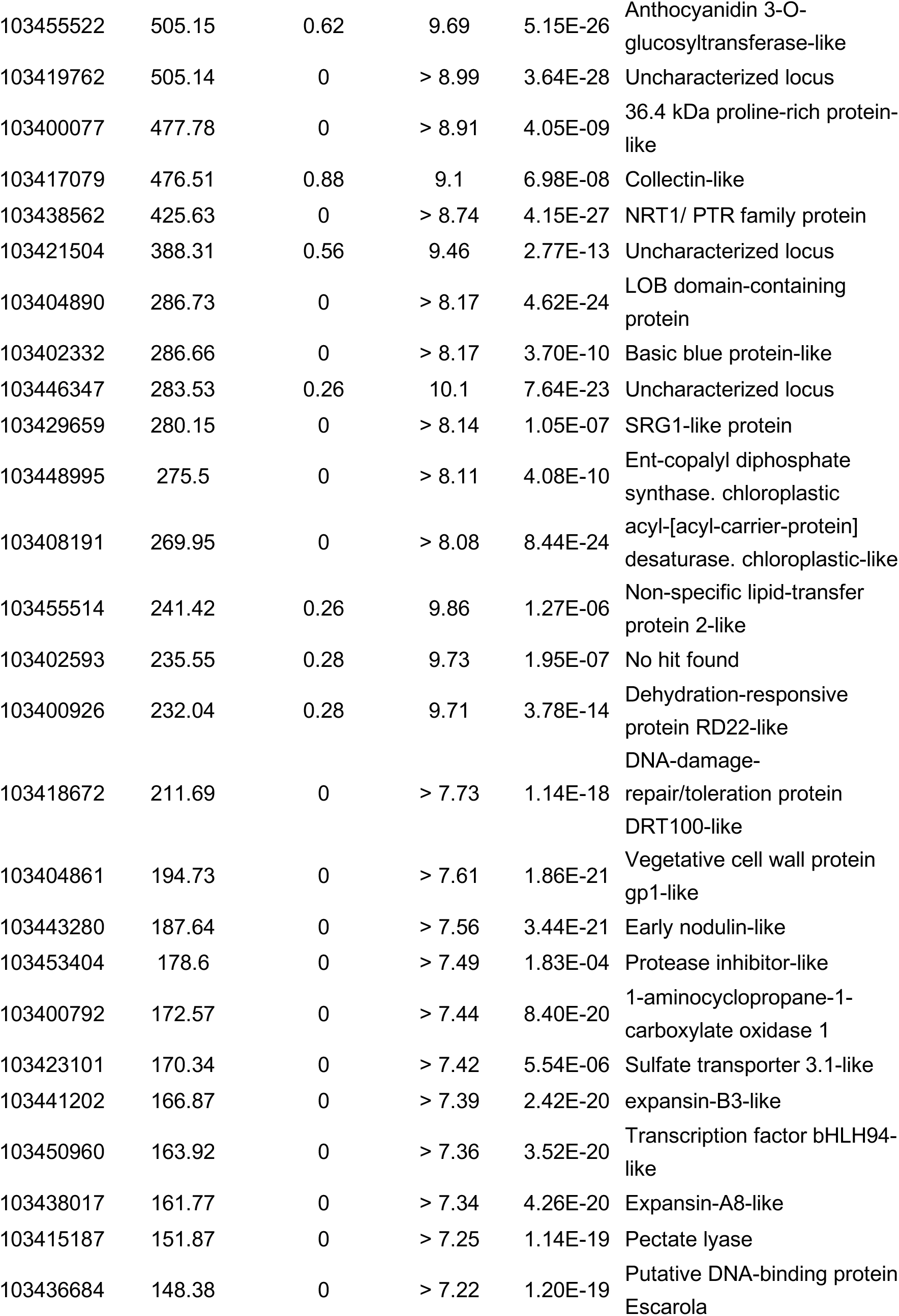
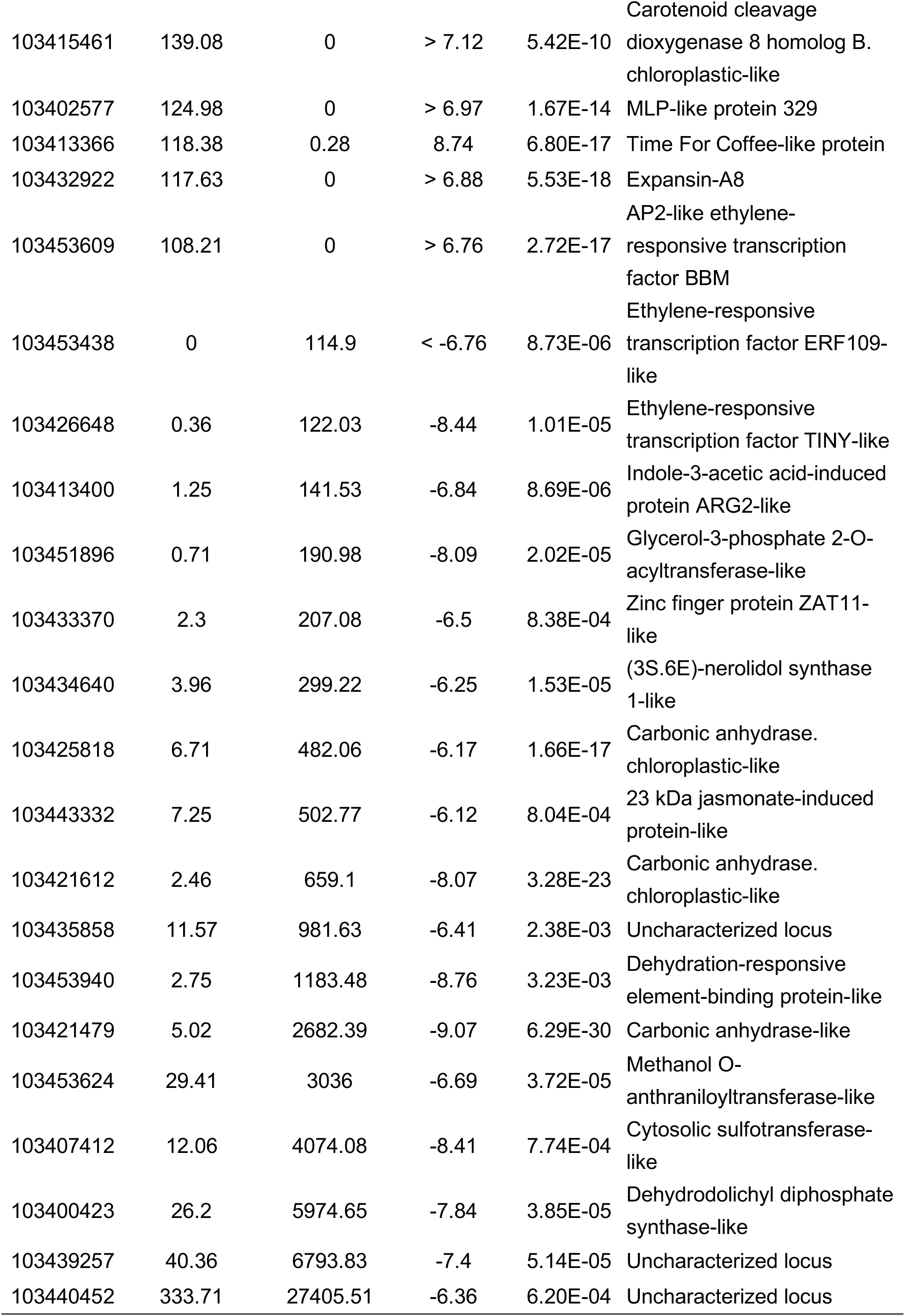
Top significantly differentially expressed apple genes (DEGs) in leaves infected by the rust fungus *Gymnosporangium yamadae* at 10 and 30 days post inoculation (dpi). Only DEGs showing a readcount over 100 in one of the conditions (inoculated or mock-inoculated) were considered at each time point.

| Gene_id | Readcount<br>Inoculated | Readcount<br>Mock-inoculated | log<sub>2</sub>(fold-change) | P-value | Gene description |
| --- | --- | --- | --- | --- | --- |
| **10 dpi** |  |  |  |  |  |
| 103443849 | 6675.97 | 21.44 | 8.29 | 3.70E-148 | Short-chain type dehydrogenase/reductase-like |
| 103415907 | 1303.09 | 3.42 | 8.58 | 3.53E-108 | Reticuline oxidase-like protein |
| 103400169 | 692.51 | 2.81 | 7.95 | 7.39E-54 | Patatin-like protein |
| 103439643 | 660.84 | 1.4 | 8.89 | 2.38E-03 | No hit found |
| 103413950 | 618.35 | 0 | > 9.28 | 3.39E-10 | Beta-glucosidase |
| 103408464 | 582.8 | 0.85 | 9.44 | 4.21E-15 | Flavanone 7-O-glucoside 2"-O-beta-L-rhamnosyltransferase-like |
| 103426327 | 452.31 | 0.85 | 9.07 | 4.27E-74 | 1-aminocyclopropane-1-carboxylate synthase-like |
| 103440114 | 349.25 | 0.87 | 8.66 | 1.54E-63 | Sorbitol dehydrogenase-like |
| 103429696 | 334.68 | 0.57 | 9.21 | 7.93E-12 | Putative 12-oxophytodienoate reductase |
| 103423646 | 318.52 | 0.29 | 10.15 | 2.25E-09 | Beta-glucosidase |
| 103404862 | 280.77 | 1.14 | 7.96 | 4.40E-16 | Formin-like protein |
| 103436320 | 251.41 | 0 | > 7.98 | 3.73E-03 | Beta-glucosidase |
| 103421201 | 245.17 | 0.86 | 8.16 | 8.61E-51 | Patatin-like protein |
| 103425667 | 220.69 | 0.57 | 8.62 | 6.93E-28 | Uncharacterized locus |
| 103447903 | 211.32 | 0.86 | 7.96 | 5.16E-30 | Bifunctional monodehydroascorbate reductase and carbonic anhydrase nectarin-like |
| 103420873 | 194.51 | 0.59 | 8.39 | 1.07E-17 | Probable LRR receptor-like serine/threonine-protein kinase |
| 103423647 | 189.96 | 0 | > 7.57 | 1.09E-02 | Cyanogenic beta-glucosidase-like |
| 103401439 | 141.45 | 0.29 | 8.98 | 2.50E-07 | Uncharacterized locus |
| 103413952 | 116.62 | 0 | > 6.87 | 1.85E-07 | Uncharacterized locus |
| 103404861 | 110.04 | 0.28 | 8.64 | 4.08E-30 | Vegetative cell wall protein<br>gp1-like |
| 103449066 | 2.29 | 112.76 | -5.63 | 5.85E-03 | Transcription initiation factor<br>TFIID subunit |
| 103440578 | 2.46 | 117.2 | -5.58 | 5.21E-03 | Aspartic Protease in Guard<br>Cell-like protein |
| 103403477 | 2.78 | 144.21 | -5.7 | 4.65E-10 | Serine/threonine-protein<br>kinase BRI1-like |
| 103440896 | 0.39 | 235.97 | -9.26 | 2.41E-11 | Thermospermine synthase<br>Acaulis-like |
| 103455791 | 2.91 | 292 | -6.66 | 2.02E-06 | Thermospermine synthase<br>Acaulis-like |
| 103437991 | 2.51 | 327.42 | -7.03 | 7.56E-12 | Zinc finger protein Constans-<br>like |
| 103451980 | 5.91 | 422.22 | -6.17 | 3.42E-05 | Galactinol synthase-like |
| 103451601 | 1.4 | 634.37 | -8.84 | 1.92E-07 | Beta-amyrin 28-oxidase-like |
| 103424346 | 22.24 | 2625.42 | -6.89 | 1.77E-04 | Uncharacterized locus |
| 30 dpi |  |  |  |  |  |
| 103451807 | 4251.52 | 0.34 | 13.64 | 1.72E-41 | Expansin-A10 |
| 103452933 | 2805.97 | 1.36 | 11.02 | 1.16E-35 | SRG1-like protein |
| 103433546 | 2182.84 | 5.02 | 8.77 | 7.86E-06 | laccase-like |
| 103425476 | 2098.2 | 0.54 | 11.94 | 8.04E-36 | Expansin-A1-like |
| 103433527 | 1359.8 | 0.95 | 10.49 | 4.09E-32 | Non-specific lipid-transfer<br>protein |
| 103438035 | 1276.53 | 0.28 | 12.17 | 3.03E-33 | Expansin-A8-like |
| 103411433 | 1080.33 | 0 | > 10.08 | 8.71E-20 | Uncharacterized locus |
| 103456171 | 970.52 | 1.32 | 9.53 | 1.05E-05 | Uncharacterized locus |
| 103441305 | 904.03 | 0 | > 9.83 | 8.59E-14 | Carotenoid cleavage<br>dioxygenase 8 homolog B.<br>chloroplastic |
| 103443282 | 835.31 | 1.71 | 8.94 | 5.04E-27 | 36.4 kDa proline-rich protein-like |
| 103441585 | 702.17 | 1.4 | 8.98 | 2.32E-26 | extensin |
| 103443566 | 686.29 | 0.8 | 9.75 | 1.78E-27 | Anthranilate N-benzoyltransferase protein |
| 103404862 | 681.53 | 0.56 | 10.27 | 3.60E-28 | Formin-like protein |
| 103452268 | 531.22 | 0 | > 9.06 | 5.88E-20 | early nodulin-93-like |
| 103402951 | 517.95 | 0.8 | 9.35 | 1.21E-25 | Transcription factor bHLH79-like |
| 103455522 | 505.15 | 0.62 | 9.69 | 5.15E-26 | Anthocyanidin 3-O-glucosyltransferase-like |
| 103419762 | 505.14 | 0 | > 8.99 | 3.64E-28 | Uncharacterized locus |
| 103400077 | 477.78 | 0 | > 8.91 | 4.05E-09 | 36.4 kDa proline-rich protein-like |
| 103417079 | 476.51 | 0.88 | 9.1 | 6.98E-08 | Collectin-like |
| 103438562 | 425.63 | 0 | > 8.74 | 4.15E-27 | NRT1/ PTR family protein |
| 103421504 | 388.31 | 0.56 | 9.46 | 2.77E-13 | Uncharacterized locus |
| 103404890 | 286.73 | 0 | > 8.17 | 4.62E-24 | LOB domain-containing protein |
| 103402332 | 286.66 | 0 | > 8.17 | 3.70E-10 | Basic blue protein-like |
| 103446347 | 283.53 | 0.26 | 10.1 | 7.64E-23 | Uncharacterized locus |
| 103429659 | 280.15 | 0 | > 8.14 | 1.05E-07 | SRG1-like protein |
| 103448995 | 275.5 | 0 | > 8.11 | 4.08E-10 | Ent-copalyl diphosphate synthase. chloroplastic |
| 103408191 | 269.95 | 0 | > 8.08 | 8.44E-24 | acyl-[acyl-carrier-protein] desaturase. chloroplastic-like |
| 103455514 | 241.42 | 0.26 | 9.86 | 1.27E-06 | Non-specific lipid-transfer protein 2-like |
| 103402593 | 235.55 | 0.28 | 9.73 | 1.95E-07 | No hit found |
| 103400926 | 232.04 | 0.28 | 9.71 | 3.78E-14 | Dehydration-responsive protein RD22-like |
| 103418672 | 211.69 | 0 | > 7.73 | 1.14E-18 | DNA-damage-repair/toleration protein DRT100-like |
| 103404861 | 194.73 | 0 | > 7.61 | 1.86E-21 | Vegetative cell wall protein gp1-like |
| 103443280 | 187.64 | 0 | > 7.56 | 3.44E-21 | Early nodulin-like |
| 103453404 | 178.6 | 0 | > 7.49 | 1.83E-04 | Protease inhibitor-like |
| 103400792 | 172.57 | 0 | > 7.44 | 8.40E-20 | 1-aminocyclopropane-1-carboxylate oxidase 1 |
| 103423101 | 170.34 | 0 | > 7.42 | 5.54E-06 | Sulfate transporter 3.1-like |
| 103441202 | 166.87 | 0 | > 7.39 | 2.42E-20 | expansin-B3-like |
| 103450960 | 163.92 | 0 | > 7.36 | 3.52E-20 | Transcription factor bHLH94-like |
| 103438017 | 161.77 | 0 | > 7.34 | 4.26E-20 | Expansin-A8-like |
| 103415187 | 151.87 | 0 | > 7.25 | 1.14E-19 | Pectate lyase |
| 103436684 | 148.38 | 0 | > 7.22 | 1.20E-19 | Putative DNA-binding protein Escarola |
| 103415461 | 139.08 | 0 | > 7.12 | 5.42E-10 | Carotenoid cleavage dioxygenase 8 homolog B. chloroplastic-like |
| 103402577 | 124.98 | 0 | > 6.97 | 1.67E-14 | MLP-like protein 329 |
| 103413366 | 118.38 | 0.28 | 8.74 | 6.80E-17 | Time For Coffee-like protein |
| 103432922 | 117.63 | 0 | > 6.88 | 5.53E-18 | Expansin-A8 |
| 103453609 | 108.21 | 0 | > 6.76 | 2.72E-17 | AP2-like ethylene-responsive transcription factor BBM |
| 103453438 | 0 | 114.9 | < -6.76 | 8.73E-06 | Ethylene-responsive transcription factor ERF109-like |
| 103426648 | 0.36 | 122.03 | -8.44 | 1.01E-05 | Ethylene-responsive transcription factor TINY-like |
| 103413400 | 1.25 | 141.53 | -6.84 | 8.69E-06 | Indole-3-acetic acid-induced protein ARG2-like |
| 103451896 | 0.71 | 190.98 | -8.09 | 2.02E-05 | Glycerol-3-phosphate 2-O-acyltransferase-like |
| 103433370 | 2.3 | 207.08 | -6.5 | 8.38E-04 | Zinc finger protein ZAT11-like |
| 103434640 | 3.96 | 299.22 | -6.25 | 1.53E-05 | (3S.6E)-nerolidol synthase 1-like |
| 103425818 | 6.71 | 482.06 | -6.17 | 1.66E-17 | Carbonic anhydrase. chloroplastic-like |
| 103443332 | 7.25 | 502.77 | -6.12 | 8.04E-04 | 23 kDa jasmonate-induced protein-like |
| 103421612 | 2.46 | 659.1 | -8.07 | 3.28E-23 | Carbonic anhydrase. chloroplastic-like |
| 103435858 | 11.57 | 981.63 | -6.41 | 2.38E-03 | Uncharacterized locus |
| 103453940 | 2.75 | 1183.48 | -8.76 | 3.23E-03 | Dehydration-responsive element-binding protein-like |
| 103421479 | 5.02 | 2682.39 | -9.07 | 6.29E-30 | Carbonic anhydrase-like |
| 103453624 | 29.41 | 3036 | -6.69 | 3.72E-05 | Methanol O-anthraniloyltransferase-like |
| 103407412 | 12.06 | 4074.08 | -8.41 | 7.74E-04 | Cytosolic sulfotransferase-like |
| 103400423 | 26.2 | 5974.65 | -7.84 | 3.85E-05 | Dehydrodolichyl diphosphate synthase-like |
| 103439257 | 40.36 | 6793.83 | -7.4 | 5.14E-05 | Uncharacterized locus |
| 103440452 | 333.71 | 27405.51 | -6.36 | 6.20E-04 | Uncharacterized locus |

**Figure 3.**
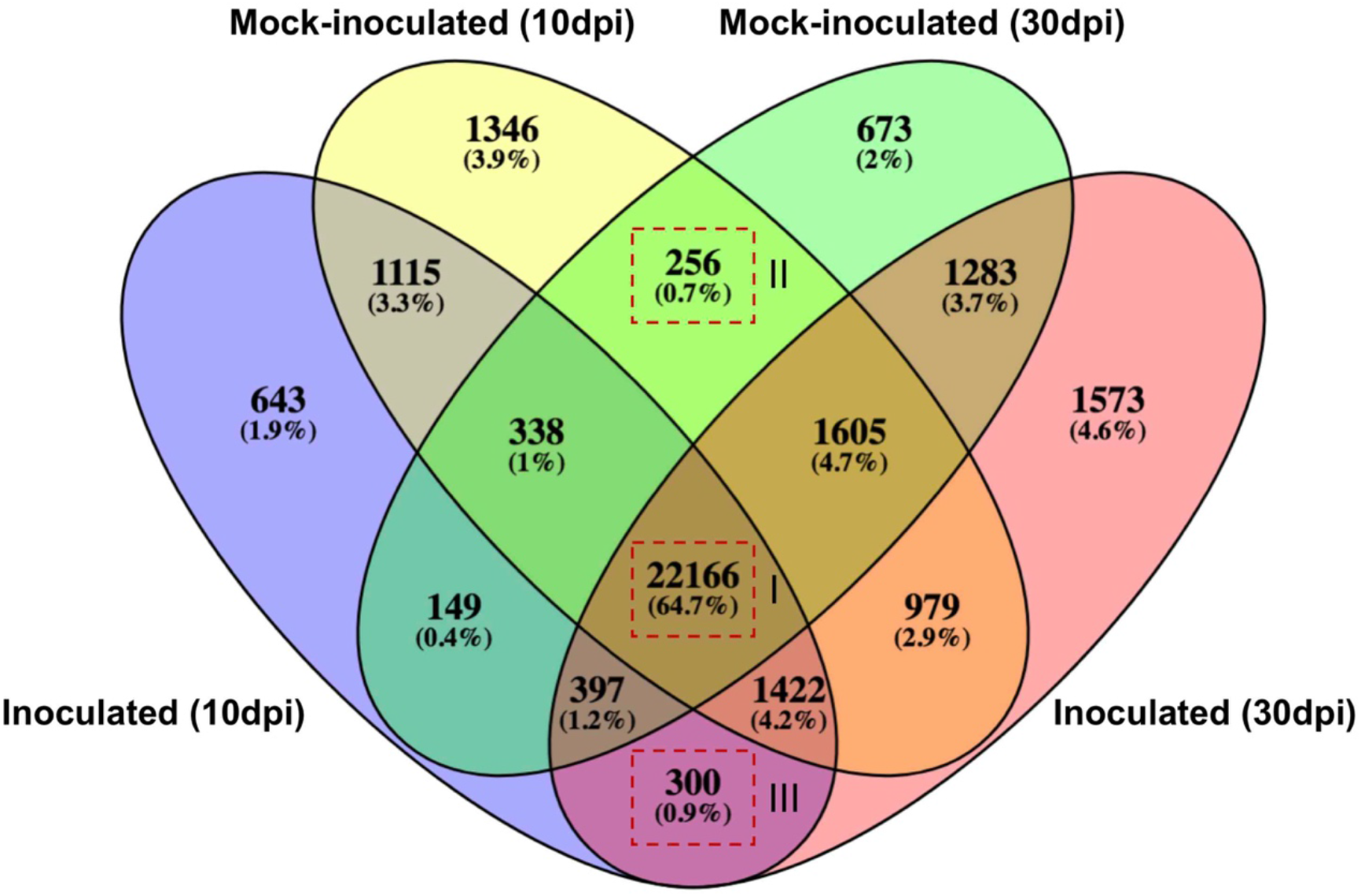
Venn diagram showing the number of genes expressed in each condition. Group I indicating the genes detected in all conditions; group II refers to genes specifically found in the inoculated groups at the two time points, 10 and 30 days post-inoculation (dpi); group III refers to genes exclusively expressed in the mock inoculated groups at two time points.

**Figure 4.**
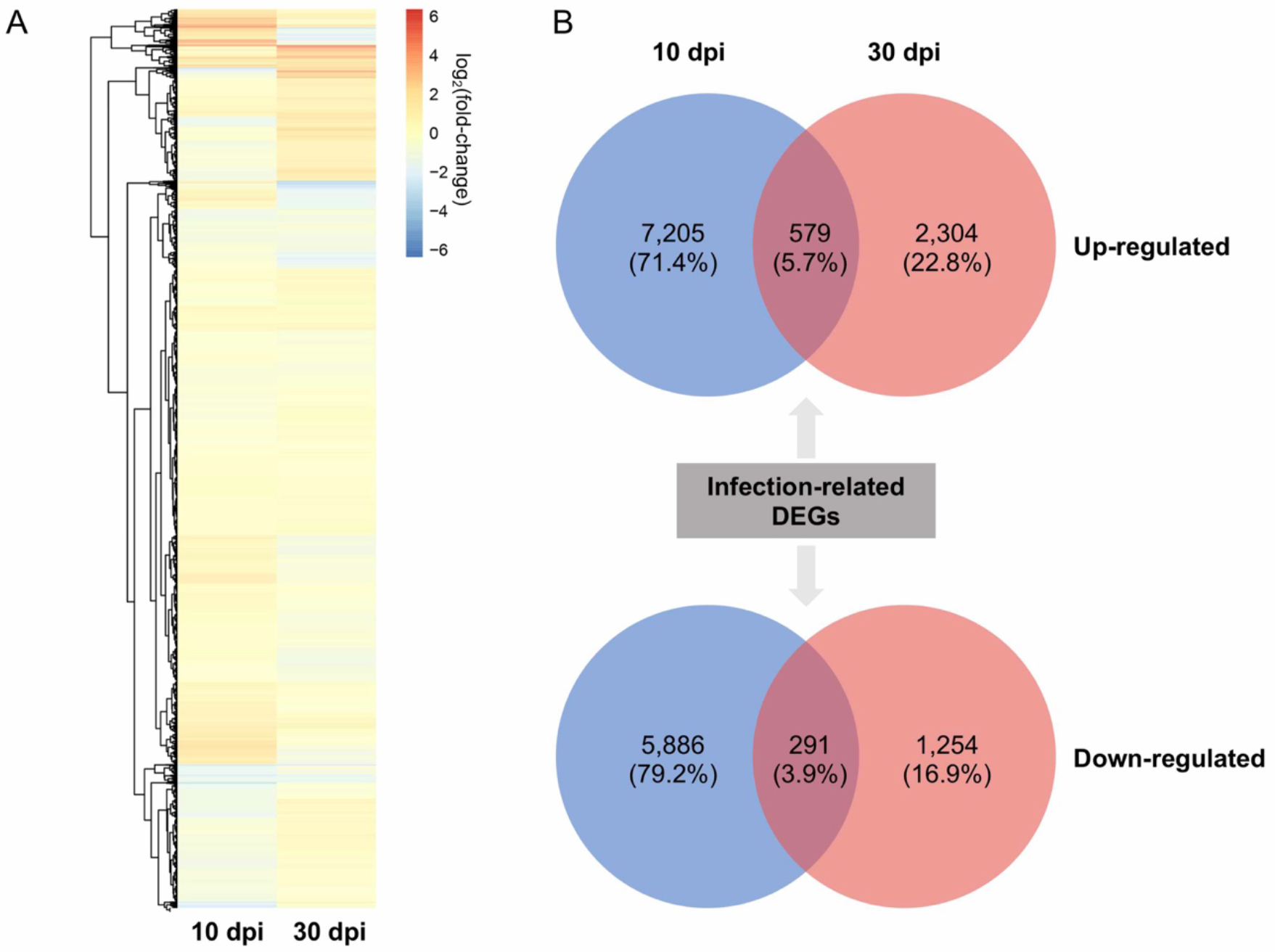
Differentially expressed genes (DEGs) between inoculated and mock inoculated apple leaves at 10 and 30 days post-inoculation (dpi). A: Hierarchical clustering analysis of the DEGs at 10 dpi and 30 dpi. B: The upper and lower Venn diagrams showing all the up-regulated and down-regulated genes at 10 dpi and 30 dpi, and the arrows pointing to the number of genes differentially expressed at the both time points.

### Impact of rust infection on metabolic pathways in apple leaves

In order to determine the impact of the rust infection on metabolic pathways of apple leaves, we performed classification of DEGs into cellular categories using MapMan and KEGG and performed enrichment analysis. All DEGs were integrated into MapMan plant categories and 52 and 56% of them were classified into 28 functional categories at 10 and 30 dpi, respectively (Figure 5A; Supplementary file: Table S1). The direct comparison of proportions of up- and down-regulated transcripts assigned to Mapman categories helped to draw major inflexion in metabolic pathways. For instance, photosynthesis processes were remarkably represented among the down-regulated transcripts at the two time points. Conversely, cellular respiration was more represented among induced transcripts at both time points in infected apple leaves (Supplementary file: Table S1, Figure 5A). Interestingly, the solute transport category was proportionally more abundant in both up- and down-regulated transcripts at 10dpi, indicating that this process might be particularly dynamic at the spermogonia infection stage. Similarly, the RNA trafficking category was also more represented in up- and down-regulated transcripts at 30 dpi, suggesting a highly dynamic regulation of this process at the aecial stage. The phytohormones and cell wall categories were also more represented among transcripts up-regulated at 10 dpi, and the vesicle trafficking category was more prominent at 30dpi among up-regulated transcripts (Figure 5A). Almost all the genes falling in the MapMan photosynthesis-related apparatus category (Photosystem I, Photosystem II, ATP synthase, Redox chain, Photorespiration and Calvin cycle) were significantly down-regulated in infected leaves at the two time points (Figure 5B).

**Figure 5.**
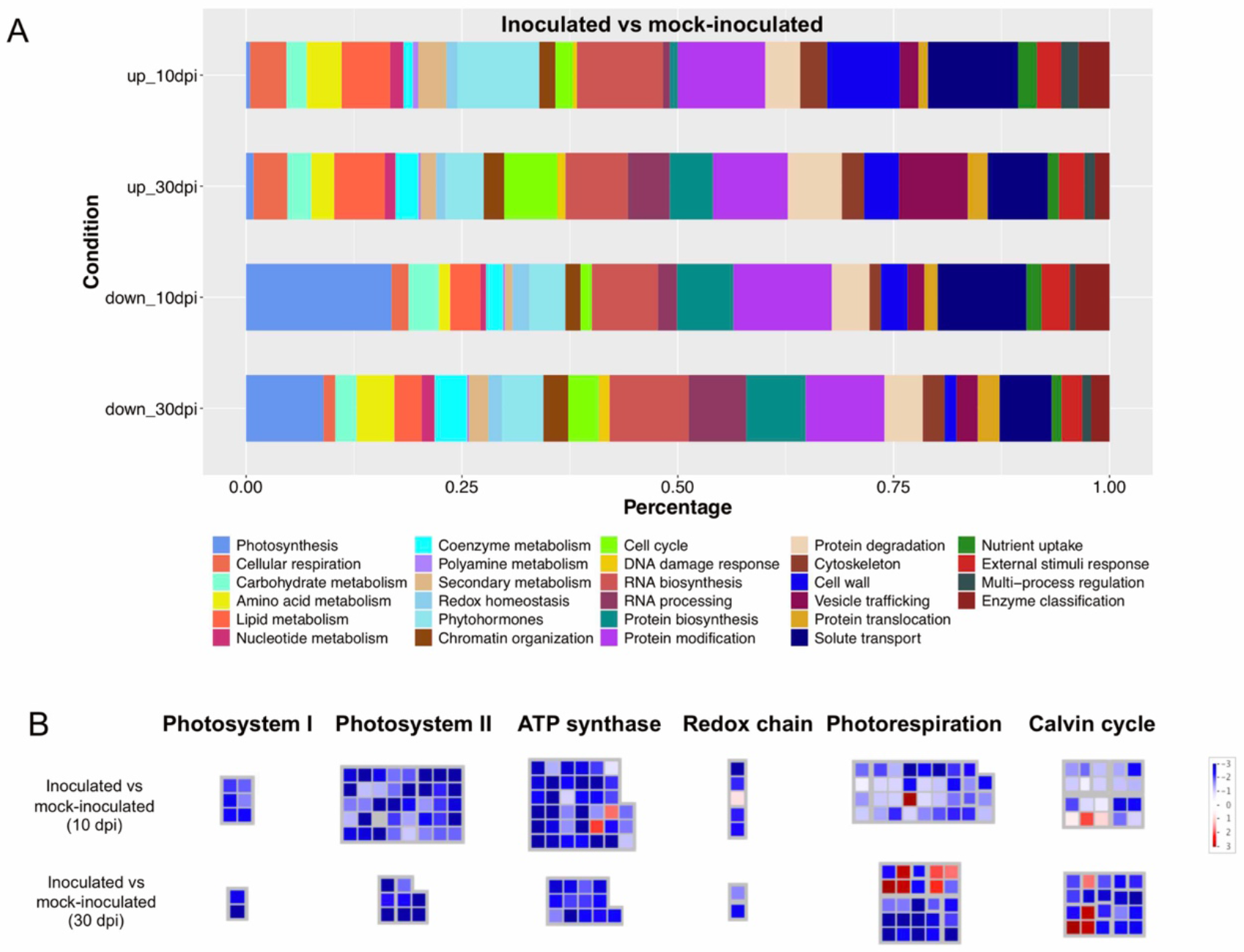
Overview of the transcriptional changes of *G. yamadae* inoculated apple leaves. A: All the differentially expressed genes at 10 and 30 days post-inoculation (dpi) were classified into MapMan metabolic pathways. The chart shows the percentage of up- or down- regulated genes at 10 dpi and 30 dpi classified into each pathway. B: Genes involved in the photosynthesis are significantly down-regulated in inoculated groups compared to mock inoculated groups at 10 dpi and 30 dpi. The photosynthetic apparatus related genes were defined according to MapMan bin code and the expression profiles were visualized by MapMan tool.

All DEGs were also classified into Kyoto Encyclopedia of Genes and Genomes (KEGG) pathways and an enrichment analysis between inoculated and mock inoculated groups was performed for down- and up-regulated transcripts. In total, 3,090 and 715 down-regulated transcripts were classified into 124 and 85 KEGG pathways, at 10 dpi and 30 dpi respectively; whereas 4,133 and 1,493 up-regulated transcripts were classified in 122 and 112 KEGG pathways at 10 and 30 dpi, respectively (Supplementary file: Table S1). Consistent with the MapMan annotation results, photosynthesis related pathways were among the most significant down-regulated pathways at both time points, including biosynthesis of light-harvesting chlorophyll protein complex (LHC) in antenna proteins, impaired processes of Photosystem I, Photosystem II, and Cytochrome b6/f complex (Supplementary file: Table S1). Glutathione metabolism, endocytosis and amino sugar and nucleotide sugar metabolism categories were significantly enriched among up-regulated transcripts at 10 dpi (Figure 6). At 30 dpi, the three most significantly enriched pathways were flavonoid biosynthesis; stilbenoid, diarylheptanoid and gingerol biosynthesis; and phenylpropanoid biosynthesis (Figure 6). Interestingly, many other pathways directly related to plant secondary metabolism such as phenylalanine metabolism or biosynthesis of secondary metabolites, were found among significantly enriched KEGG categories at this time points. Categories related to aromatic and volatile compounds or to tyrosine metabolism and tyrosine-derived alkaloids were also significantly enriched at 30 dpi. Altogether, these results point out important modifications in leaf composition. Interestingly, the plant-pathogen interaction pathway was significantly induced during infection at 10 dpi but repressed at 30 dpi (Figure 6; Supplementary file: Table S1). Overall, the survey of functional categories expressed in apple leaves in response to *G. yamadae* infection indicates that even if a compatible interaction is established by the rust fungus during its biotrophic growth, a remarkable reaction is noticeable in the leaf tissues on the host side, particularly with the expression of functions related to secondary metabolism and plant-pathogen reactions, as early as 10 dpi, and more marked by 30 dpi in inoculated compared to mock-inoculated conditions.

**Figure 6.**
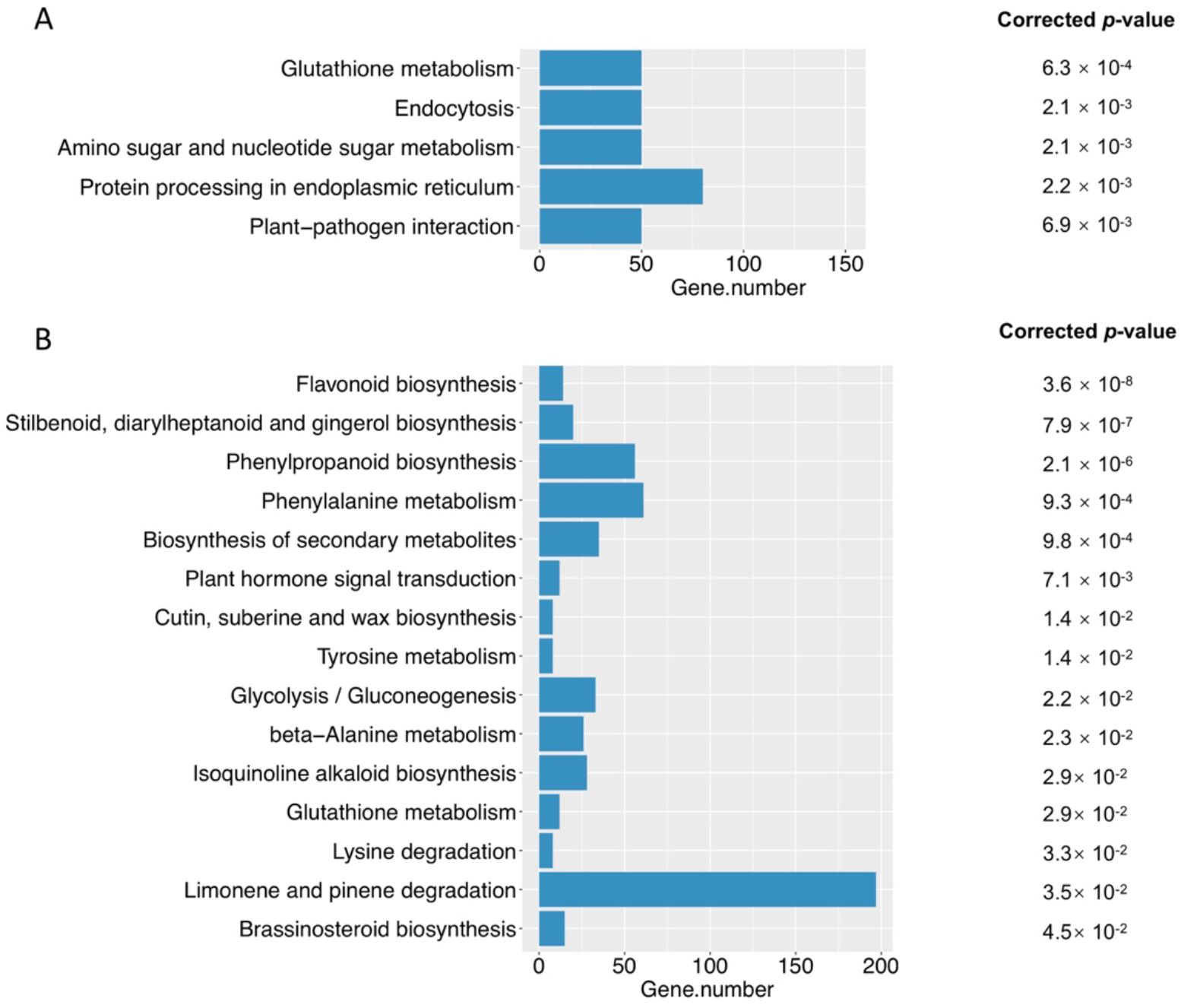
The Kyoto Encyclopedia of Genes and Genomes (KEGG) pathway enrichment analysis of all up-regulated genes in inoculated apple leaves based on the hypergeometric test. A and B showing ten most enriched pathways in inoculated apple leaves at 10 and 30 days post-inoculation (dpi), respectively. Gene number refers to the genes annotated in each pathway and corrected *p*-value is the *p*-value after Benjamini and Hochberg correction. The pathways with corrected *p*-value < 0.05 were considered as significantly enriched.

### *G. yamadae* spermogonia and aecia expressed genes

The average length of the unigenes assigned to the rust fungus in spermogonia and aecia are respectively of 1,622-bp and 1,956-bp and the largest proportion of unigenes, with 19,589 (64.67%) and 16,867 (74.25%) at 10 and 30 dpi respectively, exhibits a size larger than 1kb, indicating the good quality of the reads assembly into fungal unigenes (Supplementary file: Figure S3). Comparison of the fungal unigenes to the NCBI-nr, Swissprot and KOG databases provided putative annotation support for 17,951 (59.3%) spermogonia unigenes and 17,515 (77.1 %) aecia unigenes. Among them, 7,647 and 9,059 unigenes showed homology to genes of the wheat leaf rust fungus *Puccinia graminis* f. sp. *tritici* and 2,072 and 2,557 unigenes showed homology to genes of the poplar rust fungus *Melampsora larici-populina*. Among the 26 and 23 most highly expressed unigenes (FPKM value > 1000) in spermogonia and aecia, respectively, more than half are hypothetical proteins or without any hit in the nr database (Table 3). Beyond that, we identified transcripts encoding secreted proteins, polyubiquitin-A and a pheromone precursor highly expressed in spermogonia, and transcripts encoding secreted proteins and a thiazole biosynthetic enzyme highly expressed in aecia (Table 3). Like in other rust fungi, many highly expressed rust transcripts are of unknown function. These molecular determinants are conserved across rust species and may play an important role in pathogenesis-related processes.

**Table 3.**
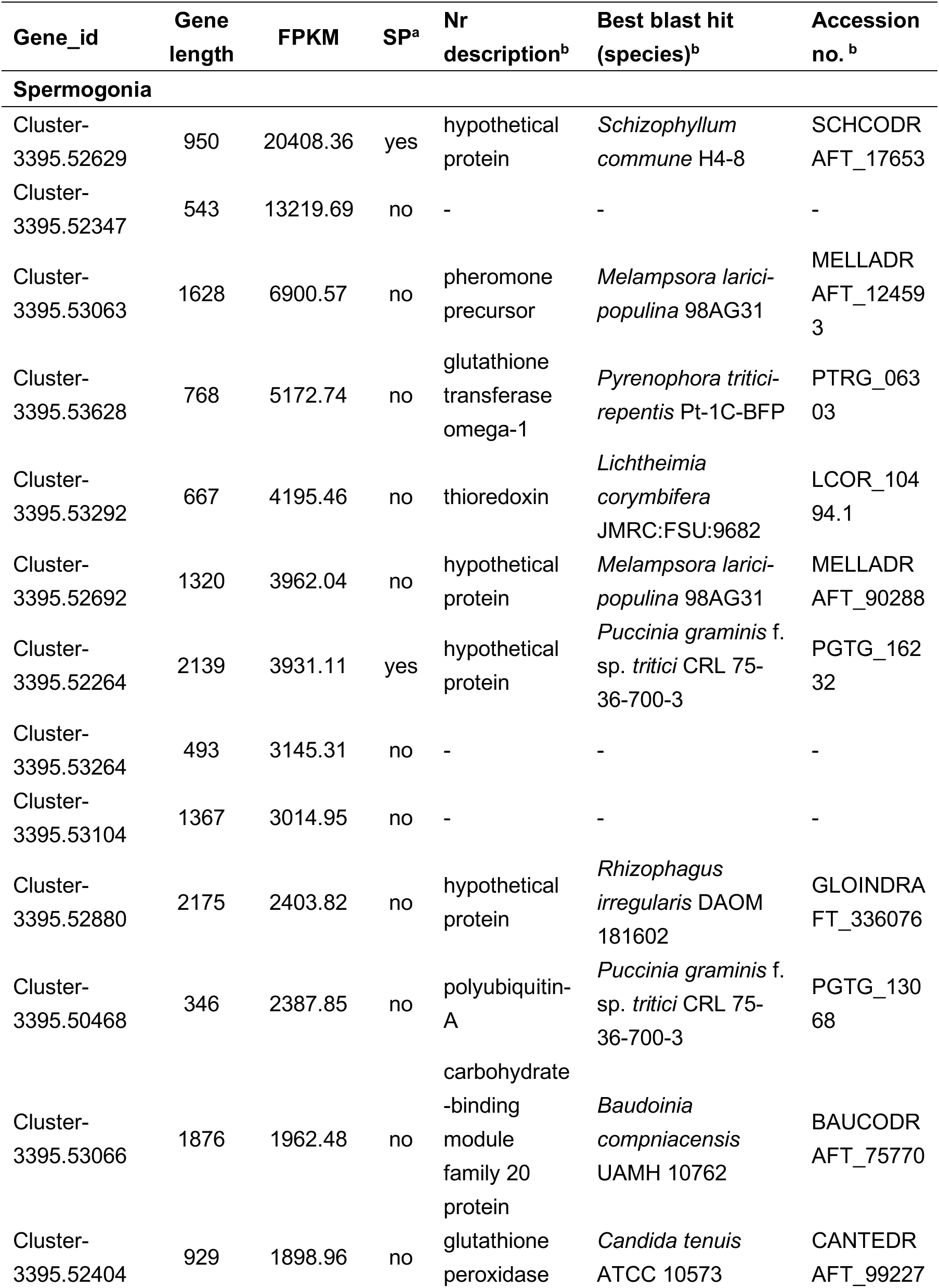
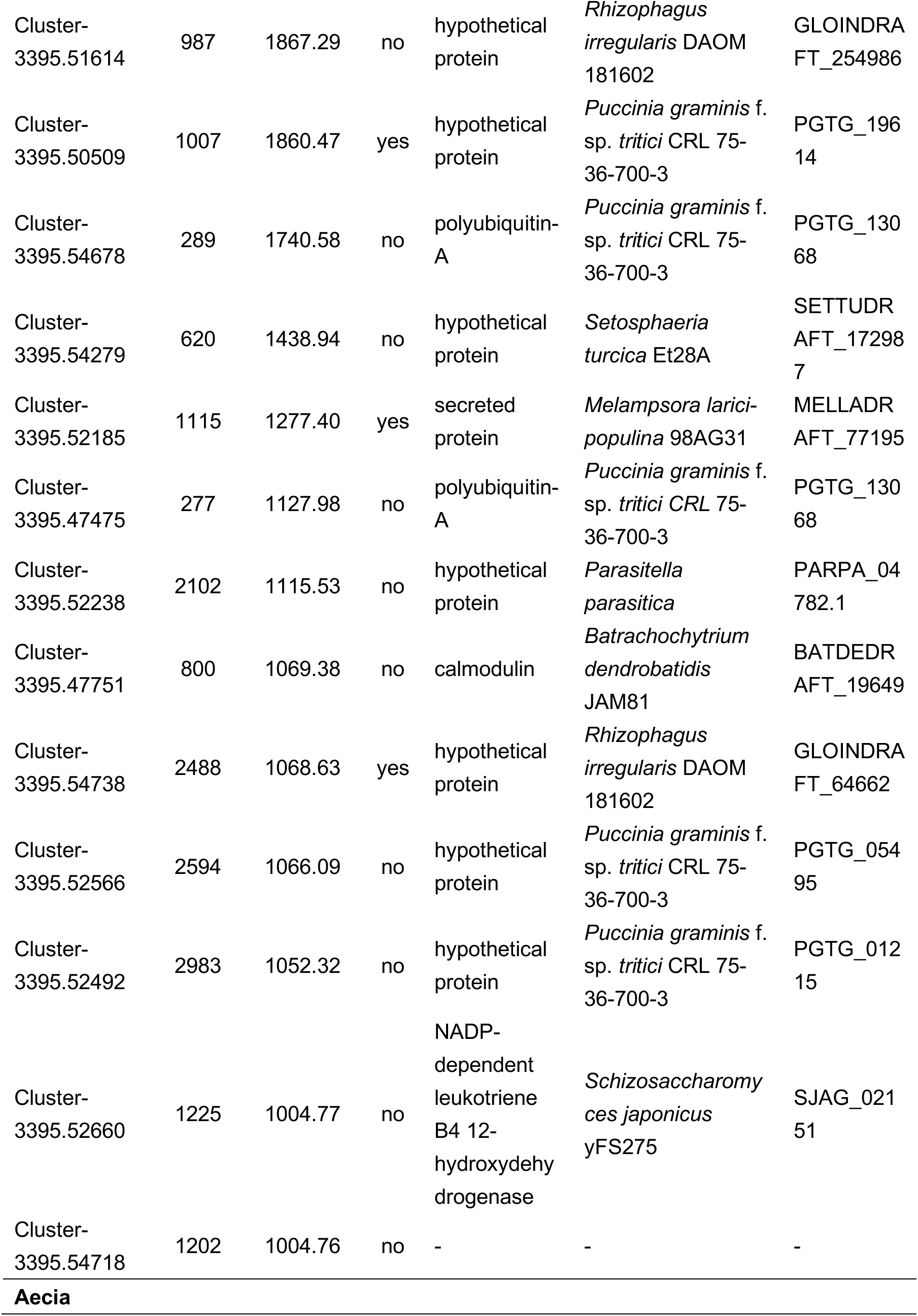
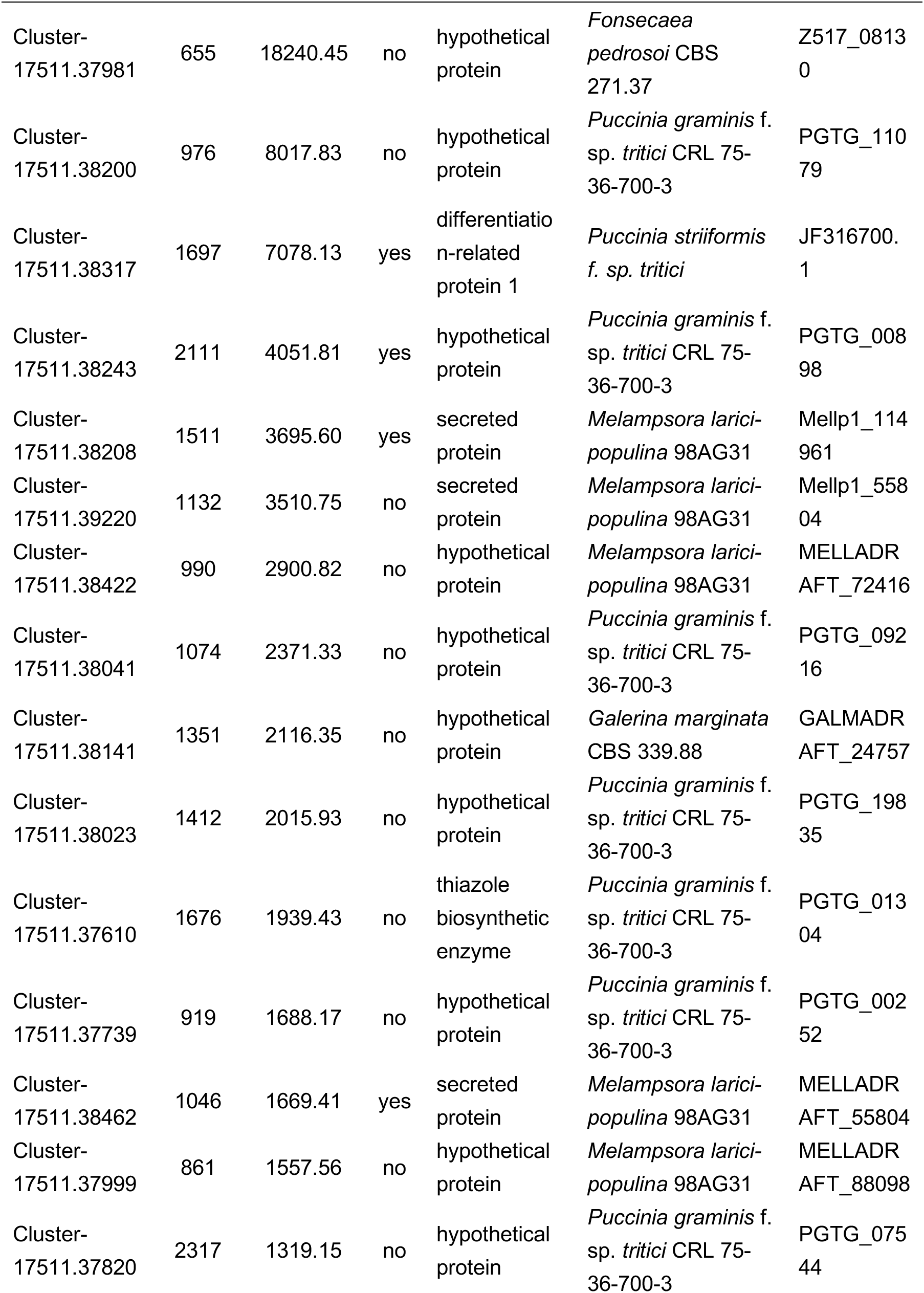
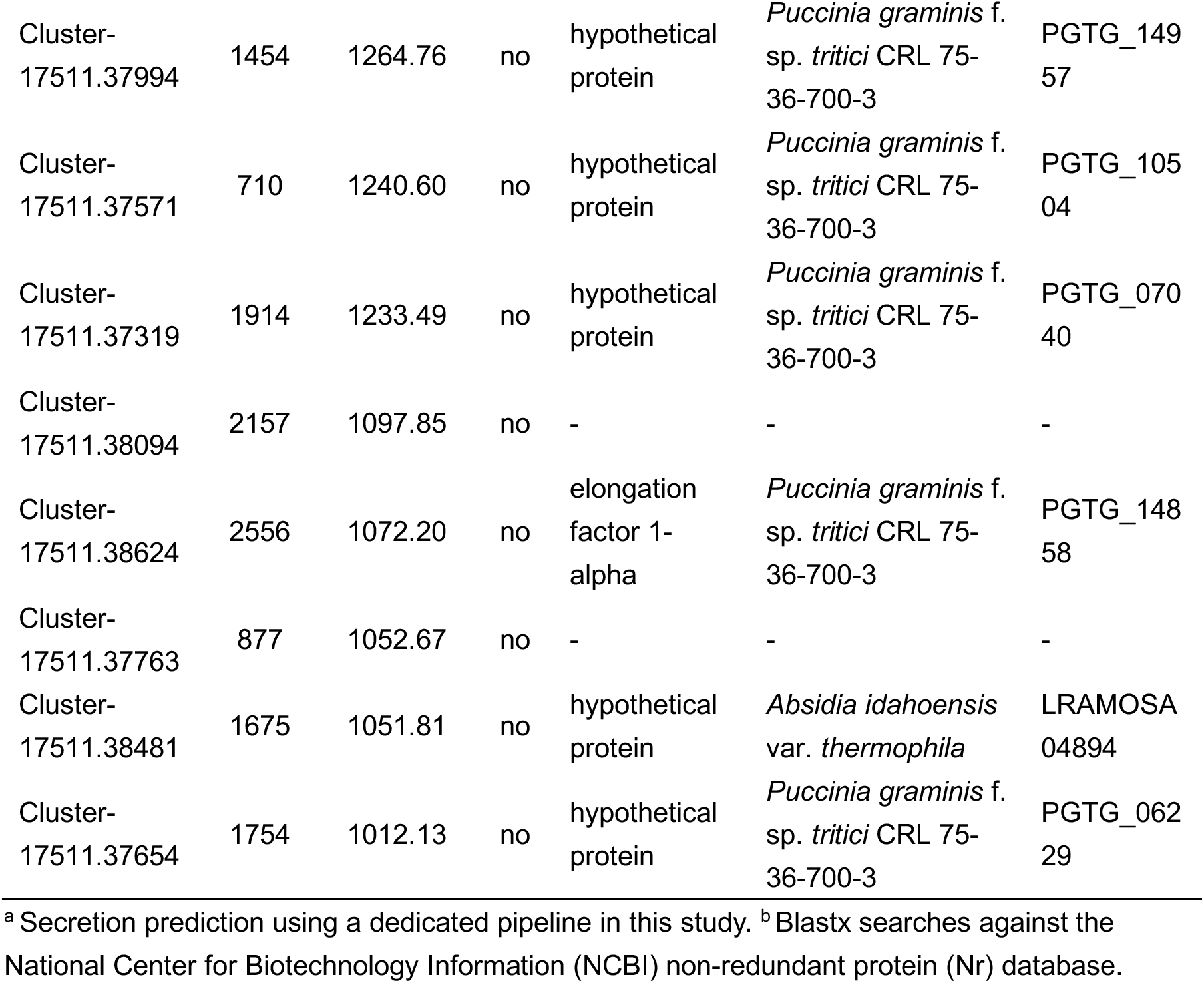
Most highly expressed *G. yamadae* unigenes (FPKM > 10^3^) in infected apple leaves at 10 days post inoculation (spermogonia) and 30 days post inoculation (aecia).

| Gene_id | Gene length | FPKM | SP<sup>a</sup> | Nr description<sup>b</sup> | Best blast hit (species)<sup>b</sup> | Accession no. <sup>b</sup> |
| --- | --- | --- | --- | --- | --- | --- |
| **Spermogonia** |  |  |  |  |  |  |
| Cluster-3395.52629 | 950 | 20408.36 | yes | hypothetical protein | *Schizophyllum commune* H4-8 | SCHCODR AFT_17653 |
| Cluster-3395.52347 | 543 | 13219.69 | no | - | - | - |
| Cluster-3395.53063 | 1628 | 6900.57 | no | pheromone precursor | *Melampsora larici-populina* 98AG31 | MELLADR AFT_124593 |
| Cluster-3395.53628 | 768 | 5172.74 | no | glutathione transferase omega-1 | *Pyrenophora tritici-repentis* Pt-1C-BFP | PTRG_06303 |
| Cluster-3395.53292 | 667 | 4195.46 | no | thioredoxin | *Lichtheimia corymbifera* JMRC:FSU:9682 | LCOR_10494.1 |
| Cluster-3395.52692 | 1320 | 3962.04 | no | hypothetical protein | *Melampsora larici-populina* 98AG31 | MELLADR AFT_90288 |
| Cluster-3395.52264 | 2139 | 3931.11 | yes | hypothetical protein | *Puccinia graminis* f. sp. *tritici* CRL 75-36-700-3 | PGTG_16232 |
| Cluster-3395.53264 | 493 | 3145.31 | no | - | - | - |
| Cluster-3395.53104 | 1367 | 3014.95 | no | - | - | - |
| Cluster-3395.52880 | 2175 | 2403.82 | no | hypothetical protein | *Rhizophagus irregularis* DAOM 181602 | GLOINDRA FT_336076 |
| Cluster-3395.50468 | 346 | 2387.85 | no | polyubiquitin-A | *Puccinia graminis* f. sp. *tritici* CRL 75-36-700-3 | PGTG_13068 |
| Cluster-3395.53066 | 1876 | 1962.48 | no | carbohydrate-binding module family 20 protein | *Baudoinia compniacensis* UAMH 10762 | BAUCODR AFT_75770 |
| Cluster-3395.52404 | 929 | 1898.96 | no | glutathione peroxidase | *Candida tenuis* ATCC 10573 | CANTEDR AFT_99227 |
| Cluster-3395.51614 | 987 | 1867.29 | no | hypothetical protein | *Rhizophagus irregularis* DAOM 181602 | GLOINDRA FT_254986 |
| Cluster-3395.50509 | 1007 | 1860.47 | yes | hypothetical protein | *Puccinia graminis* f. sp. *tritici* CRL 75-36-700-3 | PGTG_19614 |
| Cluster-3395.54678 | 289 | 1740.58 | no | polyubiquitin-A | *Puccinia graminis* f. sp. *tritici* CRL 75-36-700-3 | PGTG_13068 |
| Cluster-3395.54279 | 620 | 1438.94 | no | hypothetical protein | *Setosphaeria turcica* Et28A | SETTUDR AFT_172987 |
| Cluster-3395.52185 | 1115 | 1277.40 | yes | secreted protein | *Melampsora larici-populina* 98AG31 | MELLADR AFT_77195 |
| Cluster-3395.47475 | 277 | 1127.98 | no | polyubiquitin-A | *Puccinia graminis* f. sp. *tritici* CRL 75-36-700-3 | PGTG_13068 |
| Cluster-3395.52238 | 2102 | 1115.53 | no | hypothetical protein | *Parasitella parasitica* | PARPA_04782.1 |
| Cluster-3395.47751 | 800 | 1069.38 | no | calmodulin | *Batrachochytrium dendrobatidis* JAM81 | BATDEDR AFT_19649 |
| Cluster-3395.54738 | 2488 | 1068.63 | yes | hypothetical protein | *Rhizophagus irregularis* DAOM 181602 | GLOINDRA FT_64662 |
| Cluster-3395.52566 | 2594 | 1066.09 | no | hypothetical protein | *Puccinia graminis* f. sp. *tritici* CRL 75-36-700-3 | PGTG_05495 |
| Cluster-3395.52492 | 2983 | 1052.32 | no | hypothetical protein | *Puccinia graminis* f. sp. *tritici* CRL 75-36-700-3 | PGTG_01215 |
| Cluster-3395.52660 | 1225 | 1004.77 | no | NADP-dependent leukotriene B4 12-hydroxydehydrogenase | *Schizosaccharomyces japonicus* yFS275 | SJAG_02151 |
| Cluster-3395.54718 | 1202 | 1004.76 | no | - | - | - |
| Aecia |  |  |  |  |  |  |
| Cluster-17511.37981 | 655 | 18240.45 | no | hypothetical protein | *Fonsecaea pedrosoi* CBS 271.37 | Z517_08130 |
| Cluster-17511.38200 | 976 | 8017.83 | no | hypothetical protein | *Puccinia graminis* f. sp. *tritici* CRL 75-36-700-3 | PGTG_11079 |
| Cluster-17511.38317 | 1697 | 7078.13 | yes | differentiation-related protein 1 | *Puccinia striiformis* f. sp. *tritici* | JF316700.1 |
| Cluster-17511.38243 | 2111 | 4051.81 | yes | hypothetical protein | *Puccinia graminis* f. sp. *tritici* CRL 75-36-700-3 | PGTG_00898 |
| Cluster-17511.38208 | 1511 | 3695.60 | yes | secreted protein | *Melampsora larici-populina* 98AG31 | Mellp1_114961 |
| Cluster-17511.39220 | 1132 | 3510.75 | no | secreted protein | *Melampsora larici-populina* 98AG31 | Mellp1_55804 |
| Cluster-17511.38422 | 990 | 2900.82 | no | hypothetical protein | *Melampsora larici-populina* 98AG31 | MELLADR AFT_72416 |
| Cluster-17511.38041 | 1074 | 2371.33 | no | hypothetical protein | *Puccinia graminis* f. sp. *tritici* CRL 75-36-700-3 | PGTG_09216 |
| Cluster-17511.38141 | 1351 | 2116.35 | no | hypothetical protein | *Galerina marginata* CBS 339.88 | GALMADR AFT_24757 |
| Cluster-17511.38023 | 1412 | 2015.93 | no | hypothetical protein | *Puccinia graminis* f. sp. *tritici* CRL 75-36-700-3 | PGTG_19835 |
| Cluster-17511.37610 | 1676 | 1939.43 | no | thiazole biosynthetic enzyme | *Puccinia graminis* f. sp. *tritici* CRL 75-36-700-3 | PGTG_01304 |
| Cluster-17511.37739 | 919 | 1688.17 | no | hypothetical protein | *Puccinia graminis* f. sp. *tritici* CRL 75-36-700-3 | PGTG_00252 |
| Cluster-17511.38462 | 1046 | 1669.41 | yes | secreted protein | *Melampsora larici-populina* 98AG31 | MELLADR AFT_55804 |
| Cluster-17511.37999 | 861 | 1557.56 | no | hypothetical protein | *Melampsora larici-populina* 98AG31 | MELLADR AFT_88098 |
| Cluster-17511.37820 | 2317 | 1319.15 | no | hypothetical protein | *Puccinia graminis* f. sp. *tritici* CRL 75-36-700-3 | PGTG_07544 |
| Cluster-17511.37994 | 1454 | 1264.76 | no | hypothetical protein | *Puccinia graminis* f. sp. *tritici* CRL 75-36-700-3 | PGTG_14957 |
| Cluster-17511.37571 | 710 | 1240.60 | no | hypothetical protein | *Puccinia graminis* f. sp. *tritici* CRL 75-36-700-3 | PGTG_10504 |
| Cluster-17511.37319 | 1914 | 1233.49 | no | hypothetical protein | *Puccinia graminis* f. sp. *tritici* CRL 75-36-700-3 | PGTG_07040 |
| Cluster-17511.38094 | 2157 | 1097.85 | no | - | - | - |
| Cluster-17511.38624 | 2556 | 1072.20 | no | elongation factor 1-alpha | *Puccinia graminis* f. sp. *tritici* CRL 75-36-700-3 | PGTG_14858 |
| Cluster-17511.37763 | 877 | 1052.67 | no | - | - | - |
| Cluster-17511.38481 | 1675 | 1051.81 | no | hypothetical protein | *Absidia idahoensis* var. *thermophila* | LRAMOSA04894 |
| Cluster-17511.37654 | 1754 | 1012.13 | no | hypothetical protein | *Puccinia graminis* f. sp. *tritici* CRL 75-36-700-3 | PGTG_06229 |

<sup>a</sup> Secretion prediction using a dedicated pipeline in this study. <sup>b</sup> Blastx searches against the National Center for Biotechnology Information (NCBI) non-redundant protein (Nr) database.

### Prediction of in planta secreted proteins expressed by *G. yamadae* during apple leaf infection

Rust fungi possess very large repertoires of secreted proteins (SPs) that contain effectors, which have significant roles in the establishment of compatible interactions with their host plants (Lorrain et al. 2019). We predicted 38,039 and 29,160 proteins from *G. yamadae* spermogonia and aecia unigenes, respectively. Based on a dedicated bioinformatic pipeline, we identified 978 (2.6%) and 1,091(3.7%) secreted proteins (SPs) (Supplementary file: Table S2). In detail, the 978 spermogonia SPs contain 219 carbohydrate active-enzymes (CAZymes), 75 proteases and 11 lipases, and five of these SPs were highly expressed at this stage (FPKM value > 1000) including the transcript exhibiting the highest expression level (Cluster-3395.52629) (Supplementary file: Table S2). Among the 1,091 SPs predicted in aecia, 246, 57 and 10 proteins were classified into CAZymes, proteases and lipases, respectively. Four SPs showed a high expression levels at the aecial stage (FPKM>1000) (Supplementary file: Table S2). Predicted proteins from spermogonia and aecia were compared to proteins previously identified in *G. yamadae* telia (Tao et al. 2017) with a Markov Cluster Algorithm (MCL) analysis in order to identify proteins specific of apple infection stages (Supplementary file: Table S2). In total, 1,097 MCL protein families with a total of 3,584 proteins, only contained proteins from spermogonia and aecia (1,831 and 1,753, respectively) and are considered specific of apple infection. Besides, 2,202 MCL protein families with a total of 2,704 proteins (7.1% of 38,039 predicted proteins in spermogonia) were identified as specific of the spermogonia stage. Among these proteins, 45 were predicted as SPs. Furthermore, 1,322 protein families with a total of 1,801 proteins (6.2% of 29,160 predicted proteins in aecia) were specific to the aecia stage, including 50 SPs (Supplementary file: Table S2). Among spermogonia and aecia predicted SPs, six showed homology to the rust transferred protein 1 (RTP1), initially described in *Uromyces* spp. (Kemen et al. 2005) and since shown as conserved across all rust fungi (Lorrain et al. 2019; Pretsch et al. 2013). The six amino acid sequences ranged between 170 to 260 residues, and a sequence alignment with 13 RTP homologues from seven different rust fungi showed relatively conserved regions in the C-terminal second half of the protein with conserved cysteine positions along the sequence (Figure S4). Insterestingly, *Gymnosporangium* spp. RTP homologs clustered in two different groups, one specific of the genus with *Gymnosporangium sabinae* and another with *Hemileia vastatrix*, the coffee rust fungus, distinct from RTPs from Pucciniaceae (*Uromyces* spp.) and Melampsoraceae (Figure S4). This result indicates the presence of an ancestral and conserved rust SP multigene families for RTP in *Gymnosporangium* spp. and the expression of distinct sets of SPs -i.e. likely candidate apple rust effectors- at specific infection stages.

### Functional distribution of *G. yamadae* unigenes during the interaction with the two host plants, apple and juniper

*G. yamadae* unigenes from spermogonia and aecia were annotated in the Eukaryotic orthologous group (KOG) database. In total, 16,458 (54.33%) and 11,299 (49.74%) unigenes showed homology in the KOG database (Supplementary file: Table S2) and the functional categories were compared to the distribution previously reported for the telial stage on juniper tree (Tao et al. 2017). Excluding the category of “unknown function” and “General function prediction only”, the most abundant KOG categories corresponded to “posttranslational modification, protein turnover, chaperones” at all three stages. The KOG categories showed similar distribution across the different fungal stages, although in each KOG category the unigenes are more abundant in spermogonia and aecia. A “secreted proteins” category corresponding to SP of unknown function was included in the KOG annotation, and the genes falling into this category were much abundant in telia and aecia than in spermogonia (Supplementary file: Figure S5). Although we do not have a reference genome for *G. yamadae* and as such, we cannot accurately estimate the extent of each transcriptome completeness, this comparison suggests that similar genetic programs are expressed on the different host plants at distinct sporulation stages, with the concomitant expression of specific genes in distinct categories to achieve specific host infection and/or fungal sporulation related processes.

### Infection of *G. yamadae* modifies the composition of fungal communities in the apple phyllosphere

Unexpectedly, beyond the apple and rust transcripts, we noticed a large number of unigenes showing homology with other fungi, particularly ascomycetes. Since these sequences were also present in controlled apple leaves, they most likely correspond to resident fungal communities on or inside the leaves of the two-year-old apple seedlings used in our experimental set-up. We used a dedicated meta-transcriptomic approach to detail the fungal communities present in inoculated and in mock inoculated apple leaves at 10 and 30 dpi (Supplementary file: Figure S2). The clean reads from the 12 Illumina libraries that did not mapped onto the *M. domestica* genome were assembled together into 64,637 unigenes. Then the clean reads from each sample were assigned to these unigenes and annotated by comparison to all fungal genomes available in the MycoCosm at the Joint Genome Institute (JGI) to know the detailed fungal composition in each sample. The unigenes that did not match to MycoCosm were compared to NR at NCBI to determine their taxonomical origin. The Table S3 details the number of total and fungal unigenes found in each sample replicate. As expected, the fungal unigenes are predominant in inoculated conditions and proportionally, the fungal reads are more important at the aecial stage at 30 dpi (Figure S6). Unigenes of metazoan origin were limited in all samples and unigenes assigned to Viridiplantae still constituted the largest proportion in the control conditions and a small portion of the inoculated conditions. These sequences of plant origin that did not mapped to the apple genome may represent specific or divergent sequences from the Chinese genotype used in the study compared to the reference apple genome (Figure S6). As expected, the major part of the fungal communities in infected apple leaves are Pucciniales, with homology to *Puccinia* spp., *Melampsora* spp. and *Cronartium* spp. (Figure 7A). The average relative abundance of rust fungi is 56.5% at 10 dpi and it increases to 76.5% at 30 dpi, reflecting the increase of *G. yamadae* biomass at 30 dpi (Supplementary file: Table S3). The distribution of reads into other fungal taxonomical groups is relatively similar at all stages with the exception of the inoculated stage at 30 dpi. This different distribution is better illustrated after the removal of the reads assigned to rust fungi (Figure 7B). Unigenes with homology to *Macroventuria* spp., *Alternaria* spp., *Didymella* spp., *Ascochyta* spp. and *Boeremia* spp. from the Pleosporales order and *Lizonia* spp. were the most abundant in the mock-inoculated conditions at 10 and 30 dpi, as well as in the inoculated condition at 10 dpi. Strikingly, in the inoculated condition at 30 dpi, there is a complete shift in the fungal community composition. This aecial infection stage is marked by a large dominance of *Alternaria* spp. and *Fonsecaea* spp., whereas the other abundant ascomycetes from the Pleosporales found in the control conditions and in the inoculated condition at 10 dpi are almost absent (Figure 7B). Interestingly, a series of less represented fungal species are present in all samples and less influenced by *G. yamadae* infection (Figure 7B, Supplementary file: Table S3). This result shows that the progression of the rust disease in inoculated apple leaves strongly impacts the phyllosphere fungal community.

**Figure 7.**
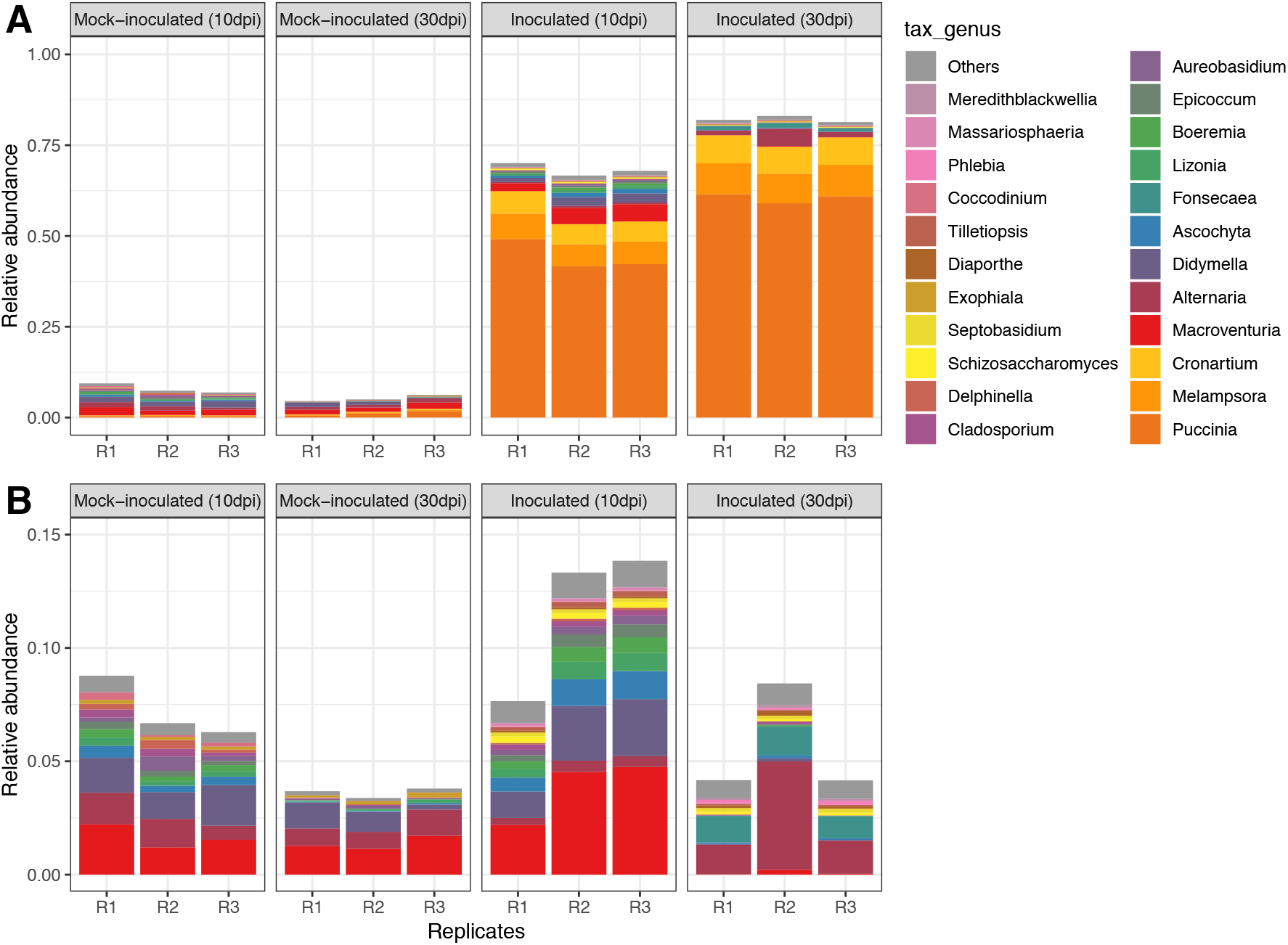
Apple phyllosphere fungal community composition in rust inoculated and mock-inoculated conditions at 10 and 30 days post-inoculation (dpi). RNA-seq reads from apple leaves unmapped to the apple genome from each biological replicate (R1 to R3) were compared to a unigenes set built from the reads of the 12 replicates altogether. The unigenes were annotated with the JGI MycoCosm genomic resource for attribution to fungal taxonomical genus (tax_genus) levels. A: relative abundance of reads attributed to unigenes annotated in fungal taxa. B: relative abundance of reads attributed to unigenes annotated in fungal taxa, after discarding rust taxa (order Pucciniales).

## Discussion

The leaf rust disease caused by the biotrophic fungus *G. yamadae* can generate substantial injuries to apple trees and result in lower fruit quality (Wang et al. 2010), and so far, there is scarce information about the molecular processes underlying the apple-apple rust interaction (Lu et al. 2017). RNA-seq has facilitated simultaneous detection of gene expression for host plants and pathogens in various pathosystems, providing new insights into understanding disease processes (Westermann et al. 2017). Availability of the apple reference genome makes it possible to monitor the transcripts expressed in infected apple leaves (Velasco et al. 2010). In the present study, we used a dual RNA-seq approach to monitor the transcriptome of apple leaves during disease progression in a compatible interaction with the rust fungus *G. yamadae* and to identify fungal genes related to pathogenesis.

Rust fungi are obligate biotrophs that establish compatible interactions to derive nutrients from their hosts (Kemen et al. 2015; Lorrain et al. 2019). In this relationship, the plant is maintained alive and the rust fungus proliferates forming specific infection structures called haustoria and later on spores that ensure propagation in the life cycle. The haustoria are formed at early stages, from which the fungus release effector proteins to interfere with the host physiology (Petre et al. 2014). From the same structures, nutrients are channelled to the fungus through specific transport systems (Struck 2015; Voegele et al. 2009). During the whole infection process, the host immunity is ineffective against the invading rust fungus, contrary to incompatible plant-rust fungus interactions that are marked by strong and early defense reactions through the specific detection of pathogen molecular determinants (Duplessis et al. 2009; Rinaldi et al. 2007). However, at late stages of compatible interactions, infected leaf tissues can present physiological reactions to rust infection during sporulation (Duplessis et al. 2009; Miranda et al. 2007; Ullah et al. 2017).

### The compatible interaction established by *G. yamadae* with apple leaves induces marked reactions including alteration of secondary metabolism pathways

Glutathione (GSH) is a tripeptide in plants playing important roles in defense reactions to pathogens, including detoxification of reactive oxygen species, regulation of formation of phytoalexin and degradation of various toxic substances through the catalyzing by GSTs (Gullner et al. 2018). In the present study, Glutathione pathway was highly enriched in inoculated apple leaves at 10 dpi according to the KEGG pathway enrichment analysis, and consistent with that, large number of GSTs encoding genes were highly induced at early infection time. Similar observations have been made during infection of poplar leaves by the rust fungus *M. larici-populina* in the frame of an incompatible interactions at early time points (2 days post-inoculation) (Rinaldi et al. 2007). Regulation of GST genes expression was also observed at later time points in compatible poplar-poplar rust interactions (Azaiez et al. 2009; Miranda et al. 2007). GSTs form a widespread and ubiquitous superfamily of plant genes, with specific expansions in trees (Lallemand et al. 2014; Pégeot et al. 2014). These enzymes have various and specific roles, some being associated to the response to rust infection (Duplessis et al. 2009; Pégeot et al. 2014). The high expression of GST genes in apple leaves may be required for detoxication of compounds accumulated during early rust infection. Beyond glutathione, the amino sugar and nucleotide sugar metabolism (ANM) pathway was also significantly enriched in infected apple leaves at 10 dpi. The products of ANM has many roles in plants, such as maintaining and repairing damaged cell walls during pathogen infection through biosynthesis of nucleotide sugar units which are components of the primary and secondary cell walls (Burton et al. 2010; Josè-Estanyol and Puigdomènech 2000; Wang et al. 2011; Wolf et al. 2012). Moreover, UPD-glucose, another important metabolite of the ANM pathway, can also be a substrate involved in callose biosynthesis (Chen and Kim 2014). Callose is widespread in higher plants and it plays important roles during plant development or in response to multiple stresses, especially to the infection of pathogens like rust fungi (Stone and Clarke 1992). The enhanced ANM metabolic pathway in *G. yamadae* infected apple leaves at 10 dpi may indicate that cell wall modification is part of the response to rust invasion at early sporulation stage during compatible interaction.

Flavonoid, phenylpropanoid and stilbenes are antimicrobial secondary metabolites known as phytoalexins that have long been associated with plant resistance (Ahuja et al. 2012). At 30 dpi, up-regulated genes were remarkably enriched in phytoalexins biosynthesis pathways, similar to the findings of the transcriptome analysis of apple leaves in response to *Marssonina coronaria* infection (Xu et al. 2015). Flavonoids are a widely distributed group of secondary metabolites in plants, including many components such as anthocyanidins, proanthocyanidins, flavanones and flavonols (Treutter 2006). The flavonoid composition positively influences the type of pigments displayed in rust infected leaves spots and the adjacent tissues, and the increased contents of flavonoid synchronized with spot expansion in apple leaves (Lu et al. 2017). Also, flavonoids biosynthesis and accumulation have been documented in poplar leaves after *Melampsora* spp. infection and biosynthesis is highly regulated by salicylic acid (Ullah et al. 2017; Ullah et al. 2019). Accumulation of proanthocyanidins and induction in expression of genes encoding enzymes involved in the synthesis of such condensed tannins were previously reported in several transcriptome analyses of compatible poplar-poplar rust interactions (Azaiez et al. 2009; Duplessis et al. 2009; Miranda et al. 2007). The over-representation of flavonoid biosynthesis pathways, secondary metabolism and pigment synthesis pathways correlates with the enlarged yellow- and red-coloured area noticeable around formation of aecia by the apple rust fungus at 30 dpi. The cumulated surface of such coloured area at the level of a single leaf can be remarkable and likely deeply alters the function of the leaves during successful disease establishment. However, it remains to be determined if this observation relates to a specific late defense reaction, for example through the loosening of the biotrophic control established by the rust fungus. It is worth to note that regulation of expression of secondary metabolism genes occurs at the uredinial sporulation stage in poplar, whereas it occurs at the aecial stage in apple leaves (Lu et al. 2017; Miranda et al. 2007; Ullah et al. 2017). No particular coloured reactions were noticeable on poplar leaves compared to apple leaves, however the time frames are different in these two different types of compatible interactions (one versus three weeks, for poplar uredinia and apple aecia, respectively). Altogether, our results and previous studies indicate that some of the plant responses to rust fungal infection are conserved independently of the stage or the type of rust life cycle (i.e. macrocyclic and demicyclic for the poplar rust and the apple rust fungi, respectively). Since the aecial stage is achieved on an alternate conifer host (larch) for the poplar rust fungus *M. larici-populina* (Lorrain et al. 2018), it would be particularly interesting to explore whether the secondary metabolic pathways show similar regulation patterns in different rust pathosystems based on plants belonging to different taxonomical groups (i.e. angiosperms versus gymnosperms).

### Transcriptome profiling of *G. yamadae* reveals expression of a conserved rust infection program and of specific in planta candidate effectors at two sporulating stages

The large genome size of *Gymnosporangium* spp. (Tavares et al. 2014) hinders sequencing in this genus. With further efforts to obtain a reference genome for this genus (Aime et al. 2017; https://jgi.doe.gov/csp-2018-duplessis-reference-genomes-50-rust-fungi/), the transcriptomes produced here and in a previous study (Tao et al. 2017) will be helpful to support genome annotation. In the absence of any supporting *G. yamadae* genome resource, a predictive annotation strategy relying on comparison to pre-existing rust genes and transcripts in databases was built after an approach described for the coffee rust fungus *H. vastatrix* (Fernandez et al. 2012), and helped the identification of fungal unigenes from infected apple leaves. The prediction-based method most likely overestimates the unigenes numbers with some redundancy in the dataset when expected gene numbers in rust fungi are considered (Aime et al. 2017). Nevertheless, the expression levels of *G. yamadae* transcripts were particularly high at apple infection stages spermogonia and aecia, compared to those recorded in telia in infected juniper host (Tao et al. 2017), although the total unigenes numbers were in a similar range at these different sporulation stages. Similar to all rust genome and transcriptome reports up-to-now, most of the transcripts expressed in spermogonia and aecia do not have functional annotation and correspond to hypothetical or unknown proteins (Aime et al. 2017).

As typical heteroecious rust fungi, *Gymnosporangium* spp. need two taxonomically unrelated hosts to complete their life cycle. The resolution of transcriptome profiles in alternate hosts has been only achieved in a few rust fungi so far (Cuomo et al. 2017; Liu et al. 2015; Lorrain et al. 2018) and the mechanisms underlying heteroecism remains largely unknown (Aime et al. 2017; Duplessis et al. 2014). Bioinformatic prediction pipelines have been used to predict rust secretomes and identify SPs representing candidate rust effectors (Lorrain et al. 2019; Sperschneider et al. 2017). The overrepresentation of specific SPs among *M. larici-populina* regulated genes during infection of its two different hosts suggests that such candidate effectors may underly establishment of compatible interactions with different host plants (Lorrain et al. 2018). In apple leaves infected by *G. yamadae*, higher proportions of SPs of unknown functions were identified in telia and aecia than in spermogonia. The expanded numbers of SPs in telia and aecia may relate to the potential roles of SPs in host alternation. *G. yamadae* SP gene families showing preferential expression during apple leaf infection in spermogonia or aecia represent priority apple rust candidate effectors for further functional characterization to understand their precise role in pathogenesis. Beyond specific *G. yamadae* SP genes, several rust-conserved SP genes were expressed during apple infection. The haustorially expressed SP RTP1 was firstly identified in *Uromyces fabae* and *U. striatus* in which it was shown to be transferred into the host cell (Kemen et al. 2005). Since, many homologues of RTP1 were found in different rust fungi, establishing a conserved rust SP gene family (Fernandez et al. 2012; Pretsch et al. 2013; Puthoff et al. 2008). Here, six unigenes retrieved in spermogonia and aecia define new members of this ancient rust SP family. The phylogeny of the RTP family shows the existence of a specific *Gymnosporangium* spp. RTP clade, which may indicate a specific evolution in this particular fungal family in the order Pucciniales.

The distribution of expressed unigenes of the apple rust fungus in functional KOG categories shows similar overall patterns in spermogonia and in aecia in apple tree, as well as in telia on the alternate host juniper. The same conclusion was reached for *M. larici-populina* on the two hosts, poplar and larch (Lorrain et al. 2018). These transcriptomes studies suggest the expression of conserved molecular mechanisms in rust fungi during infection of different hosts and at different sporulation stages. Beyond the requirement of specific initial sets of effectors to bypass the host immune system, the redundant genetic programs underlying the biotrophic growth may explain why rust fungi can successfully infect taxonomically unrelated alternate host plants in a same life cycle. A more systematic survey of transcriptomes expressed in alternate hosts in different pathosystems established with rust fungi from different taxonomical families is needed to validate this hypothesis.

### The fungal community composition of the apple phyllosphere is altered by rust infection

Plant phyllosphere represents one of the largest habitats for diverse community of prokaryotic and eukaryotic microorganisms (Lindow et al. 2003). Some resident species are plant pathogens, but most microorganisms are non-pathogenic and have been shown to play a critical role in promoting plant growth and protecting plant against pathogens (Vorholt 2012). Plant growth-promoting microbes have attracted much attention in recent years, since understanding their roles in plant could be crucial in controlling disease severity. The fundamental roles of leaf-colonizing bacteria in plant-host fitness have been analysed to a great extent and fungal colonizers in the phyllosphere, like their bacterial counterparts, form diverse communities and have been shown to modify disease severity in their host plants through interacting with pathogens or activating plant defense mechanism (Arnold et al. 2002; Busby et al. 2015; Laforest-Lapointe et al. 2019). As plants are facing environmental stresses during their growth period, microbes from the phyllosphere are also exposed to many biotic and abiotic constraints, however, the impact of pathogens on fungal communities and diversity in the phyllosphere are largely underexplored. In our study, high-throughput RNA-seq has allowed to determine the fungal communities present inside or on the surface of the apple phyllosphere and to see how they are affected by *G. yamadae* infection. The fungal communities of infected and healthy apple leaves changed between 10 dpi and 30 dpi. Such changes may be due to the variation of environmental conditions (e.g. moving from indoor to outdoor in the procedure to obtain spermogonia and aecia) which may lead to changes in leaf metabolites and further affect the growth of fungal species (Gomes et al. 2018), or it could be due to the challenge from natural microbes (Yang et al. 2016), or both. The fungal genus *Alternaria* and *Fonsecaea* presented a remarkable shift in abundance in inoculated leaf tissues at 30 dpi compared to all other conditions, including infection at 10 dpi. Since our initial experimental design was not established to specifically survey the fungal community composition in apple leaves, our report is merely descriptive and correlative. We cannot determine whether the composition change observed at 30 dpi is due to the host reaction to rust infection, or directly from the fungus biotrophic growth through challenging host immunity and/or physiology, or both. However, the effect of rust infection at 30 dpi is consistent and supported by biological replicates and it represents an interesting foundation for future studies to determine to which extent rust infection modifies the apple leaf fungal community. Many studies reported the effect of hyper-parasitic fungi on rust fungi (Kapooria and Sinha 1969; Koç and Défago 2008; Moricca et al. 2001; Tsuneda et al. 2011; Yuan et al. 1999), including one *Alternaria* species recently reported as hyper-parasites of *Puccinia striiformis* f. sp. *tritici* urediniospores (Zheng et al. 2017). The dominant position of *Alternaria* and *Fonsecaea* observed in *G. yamadae* infected leaves may imply the detection of novel *G. yamadae* hyperparasites, which may act as antagonists and represent interesting new leads for biological control of apple rust disease.

## Materials and Methods

### Preparation of plant material and artificial inoculation

Thirty *Malus domestica* cv ‘fuji’ seedlings (two-year-old, average 50 cm height) were bought from the Wanlü Nursery in Nanjing, Jiangsu, China. All these seedlings were planted in flower pots and placed in greenhouse under controlled temperature (26°C), relative humidity of 75% and with a 12h light/12h dark cycle for about one month. In early spring, *Juniperus chinensis* twigs with overwintering *G. yamadae* galls were collected in the field when mature telia extruded from the galls. The galls represent a natural inoculum, i.e. a mixture of *G. yamadae* isolates, and no other *Gymnosporangium* spp. can be confused with *G. yamadae* galls on *J. chinensis*. All these twigs were cleaned by a small brush to remove dust on the surface and placed in a container with sterile water overnight to ensure germination of telia. The germination efficiency was checked under the microscope (Leica-DM3000). Basidiospores were thoroughly mixed with sterile water and redistributed into separate sterile vials used for inoculation. Before the inoculation manipulation, all the seedlings were divided into two groups of fifteen seedlings kept under dark conditions for 12h overnight. One group was sprayed with the basidiospores solution on the top of the apple leaves and the other group was sprayed with sterile water as mock-inoculated control. After inoculation, all seedlings were covered with plastic transparent bags in order to ensure controlled infection. All bags had openings at the top to ensure air exchange. The seedlings were placed in a chamber with a temperature of 20°C and more than 95% relative humidity for two days to facilitate spore germination and penetration. Two days after inoculation, the temperature was adjusted back to room temperature (26°C) and to a relative humidity of 75%. At 10 dpi, globoid yellow spermogonia appeared on the upper surface of apple leaves. All plastic bags were then removed and the seedlings were placed outdoors for 20 more days in order to ensure proper aecia differentiation. At 30 dpi, long tubular aecia formed on the lower surface of apple leaves. At both time points, three independent replicates (different leaves from different seedlings) of approximately 50 mg diseased leaves and 50 mg control leaves were collected simultaneously. At 10 dpi, collected samples corresponded to infected leaves with bright-yellow spermogonia in the center (Figure 1). At 30 dpi, the collected samples consisted of leaf pieces with tubular aecia in the center and discoloured spots around the fungal sporulation structure. The samples were immediately frozen in liquid nitrogen and stored at −80°C until RNA isolation.

### RNA isolation, cDNA library preparation and RNA-sequencing

Apple leaves were ground to fine powder by RNase free mortars and pestles with liquid nitrogen and the total RNA were isolated using the RNeasy Plant Mini Kit (Qiagen, Beijing, China) according to the manufacturer’s instructions. For each sample, 3 μg RNA was used to generate sequencing library by using NEBNext® UltraTM RNA Library Prep Kit for Illumina® (NEB, USA). PCR was carried out using Phusion High-Fidelity DNA polymerase, universal PCR primers and Index (X) Primer. PCR products were purified by AMPure XP system (Beckman Coulter, CA, USA) and the cDNA library quality was assessed on the Agilent Bioanalyzer 2100 system (Agilent Technologies, CA, USA). The cBot Cluster Generation System was used to cluster index-coded samples in libraries by TruSeq PE Cluster Kit v3-cBot-HS (Illumina, CA, USA). In each library, 150-bp paired-end reads were generated from Illunima Hiseq platform. cDNA and RNAseq library were performed at Novogene (Beijing, China), following standard Illumina’s procedures. Raw reads in fastq format were firstly processed through in-house perl scripts to remove adapters, poly-N and low-quality reads. The remaining high-quality clean reads were used in the subsequent analysis. All raw sequence data generated in this study has been deposited in the NCBI Sequence Read Archive (https://submit.ncbi.nlm.nih.gov/subs/sra/) under the accession number SRR9326001-SRR9326012.

### Read mapping to the apple reference genome and gene expression analysis

The reference genome and gene model annotation files of *Malus domestica* v1.0 were downloaded from Phytozome v.12 (https://phytozome.jgi.doe.gov/pz/portal.html). Bowtie v2.2.3 (Langmead et al. 2019) was used to build the index of the apple reference genome and paired-end clean reads were aligned to the reference genome using Tophat v2.0.12 (Trapnell et al. 2009). The mapped reads were counted using HTSeq v0.6.1 (Anders et al. 2015) and expected number of Fragments Per Kilobase of transcript sequence per Millions base pairs sequenced (FPKM) of each gene was calculated to estimate gene expression levels. Mapping and unigene expression estimates were performed by Novogene (Beijing, China). The similarity between samples at the expression levels was visualized using R package pheatmap (https://CRAN.R-project.org/package=pheatmap) by calculating the Pearson correlation coefficient between samples. To assess the variability among samples, principle component analysis (PCA) was performed from read counts via the plotPCA function in R package DEseq2 (Love et al. 2014). The comparisons between the transcripts lists from four condition were conducted via the interactive tool Venny (http://bioinfogp.cnb.csic.es/tools/venny/index.html). Differential expression analysis between inoculated groups and mock inoculated groups was performed by DESeq2 (Love et al. 2014) using a model based on the negative binomial distribution and the resulting *p-*values were adjusted using the Benjamini and Hochberg’s approach for controlling the false discovery rate. Genes with an adjusted p-value lower than 0.05 were deemed as significantly differentially expressed.

### Functional analysis of differentially expressed genes

All the significantly differentially expressed genes between inoculated and mock-inoculated conditions were classified into MapMan functional plant categories (denominated BINs) using the automated annotation pipeline Mercator 4 with default parameters (Schwache et al. 2019). Additionally, enrichment analysis of Kyoto Encyclopedia of Genes and Genomes (KEGG) pathways was performed by KOBAS version 2.0 (Mao et al. 2005) based on the hypergeometric test, *p*-values of KEGG pathways were corrected using Benjamini and Hochberg method. Pathways with adjusted *p*-value < 0.05 were considered as significantly enriched.

### Transcriptome analysis, functional annotation, secretome prediction and sequence analysis of *G. yamadae* spermogonia and aecia

Reads unmapped to the apple reference genome in infected groups at 10d pi and 30 dpi were collectively subjected to *de novo* assembly using Trinity (Grabherr et al. 2011) with default parameters, which generated two transcriptomes at the two time points, respectively. FPKM expression levels for all unigenes were estimated in each replicate by RSEM (Li et al. 2015) and unigenes with FPKM > 0.3 in each library were retained for downstream analyses. All unigenes were compared to the genomes (blastn, e-value ≤ 10^−5^); predicted gene models (blastn, e-value ≤ 10^−5^) and predicted proteins (blastx, e-value ≤ 10^−5^) of the four rust fungi *Puccinia graminis* f. sp. *tritici*, *P. striiformis* f. sp. *tritici*, *P. triticina*, *Melampsora larici-populina* and the genome of the basidiomycete *Laccaria bicolor*, whose genomes are available in the JGI Mycocosm (Grigoriev et al. 2013). In parallel, the unigenes were compared to 97,304 cleaned Pucciniales ESTs retrieved from dbEST at GenBank (blastn, e-value ≤ 10^−5^). Unigenes showing homology in any of the searched databases were deemed as *G. yamadae* unigenes.

The fungal unigenes were annotated using public protein databases, including the National Center for Biotechnology Information (NCBI) non- redundant protein and nucleotide (NR, NT) databases, Swiss-Prot, Eukaryotic Orthologous Groups (KOG), Gene ontology (GO), protein families (PFAM), KEGG Orthology (KO) using Blastx (e-value ≤ 10^−5^). The proteomes of *G. yamadae* spermogonia and aecia stages were predicted from the corresponding unigenes using TransDecoder v3.0.1. Assignment of unigenes to rust fungi, unigene expression estimates and annotations were performed by Novogene (Beijing, China). A secretome prediction pipeline with a combination of bioinformatic tools, including SignalP v.4, WolfPSort, TMHMM, TargetP and PS-Scan algorithms, was used to predict secreted proteins as described in Pellegrin et al. (2015). Candidate CAZymes, proteases and lipases were annotated in the predicted secretomes using dbCAN v2.0 HMM-based CAZy annotation server (Zhang et al. 2018), Merops (Rawlings et al. 2016) and the Lipase Engineering database (Fischer and Pleiss 2003), respectively. The three proteomes predicted from *G.yamadae* spermogonia and aecia from this study and from *G. yamadae* telia (Tao et al. 2017) were used for comparison by MCL analysis. Gene families were clustered with fastOrtho MCL v12.135 (Wattam et al. 2014) using inflation parameters of 3 and 50% identity and coverage. Spermogonia and aecia RTP homologs were retrieved from the annotation files and aligned with selected RTP homologs from rust fungi found in NCBI with the Omega cluster (https://www.ebi.ac.uk/Tools/msa/clustalo/) (Madeira et al. 2019), and the UPGMA tree was done with MAFFT v6.864 (https://www.genome.jp/tools-bin/mafft).

### Identification of fungal community of the apple phyllosphere through a meta-transcriptomic approach

The meta-transcriptomic analysis was run separately to the transcriptome analysis using a dedicated approach and a distinct bioinformatic pipeline. First, all Illumina reads from the different replicate samples were trimmed and mapped onto the *M. domestica* reference genome using CLC Genomics Workbench 11.0 (QIAgen S.A.S. France, Courtaboeuf). A systematic trimming of 15 and 10 nucleotides at the 5’ and 3’ end, respectively, of all Illumina reads was applied, followed by mapping onto the apple genome (CLC “map reads to reference” procedure with default parameters except for length and similarity fractions set at 0.8). The final numbers of unmapped reads were in the same range than those of the transcriptome analysis. Paired-end reads unmapped onto the apple reference genome from all samples were used altogether to perform a *de novo* co-assembly using Megahit version 1.1.3 with default parameters (Li et al. 2015), and contigs smaller than 500bp were discarded. Reads of each sample were then mapped on the selected contigs using bowtie version 2.3.0 (Langmead and Salzberg 2012). Counts were determined using SAMtools version 1.7 (Li et al. 2009). Contigs supported by less than 3 samples and less than 5 counts were discarded. The remaining contigs were annotated using a blast-like procedure using DIAMOND version 0.9.19 (Buchfink et al. 2015) with the parameters --more sensitive --max-target-seqs 1 --max-hsps 1 --evalue 0.00001 and JGI-Mycocosm (https://genome.jgi.doe.gov/mycocosm/home) predicted proteins from fungal genomes (deposited before July 2018) as a reference database. Diamond annotation was doubled using NCBI-NR (March 2018 version) to check for fungal false positives, based on the comparison between Mycocosm and NR best bit scores. Count tables and annotations for phyllosphere fungal composition were then analyzed at various taxonomical levels using R version 3.4.3 and packages dplyR and ggplot2. Taxonomical category “others” corresponds to taxa with a relative abundance below 10^−3^
 in all samples.

## Acknowledgements

This work was financed by the National Natural Science Foundation of China (No. 31870628). The authors acknowledge the financial support from China Scholarship Council (No. 201806510009) for the Joint PhD Program between Beijing Forestry University and INRA-Nancy. SD is supported by the French ‘Investissements d’Avenir’ program (ANR-11-LABX-0002-01, Lab of Excellence ARBRE). Y. M. Liang and S. Duplessis designed this research. S. Q. Tao performed the experiments. L. Auer and E. Morin performed the bioinformatic analysis. S. Duplessis and S. Q. Tao analysed data and wrote the manuscript. All authors approved the final manuscript for submission.

---

**Figure S1.**
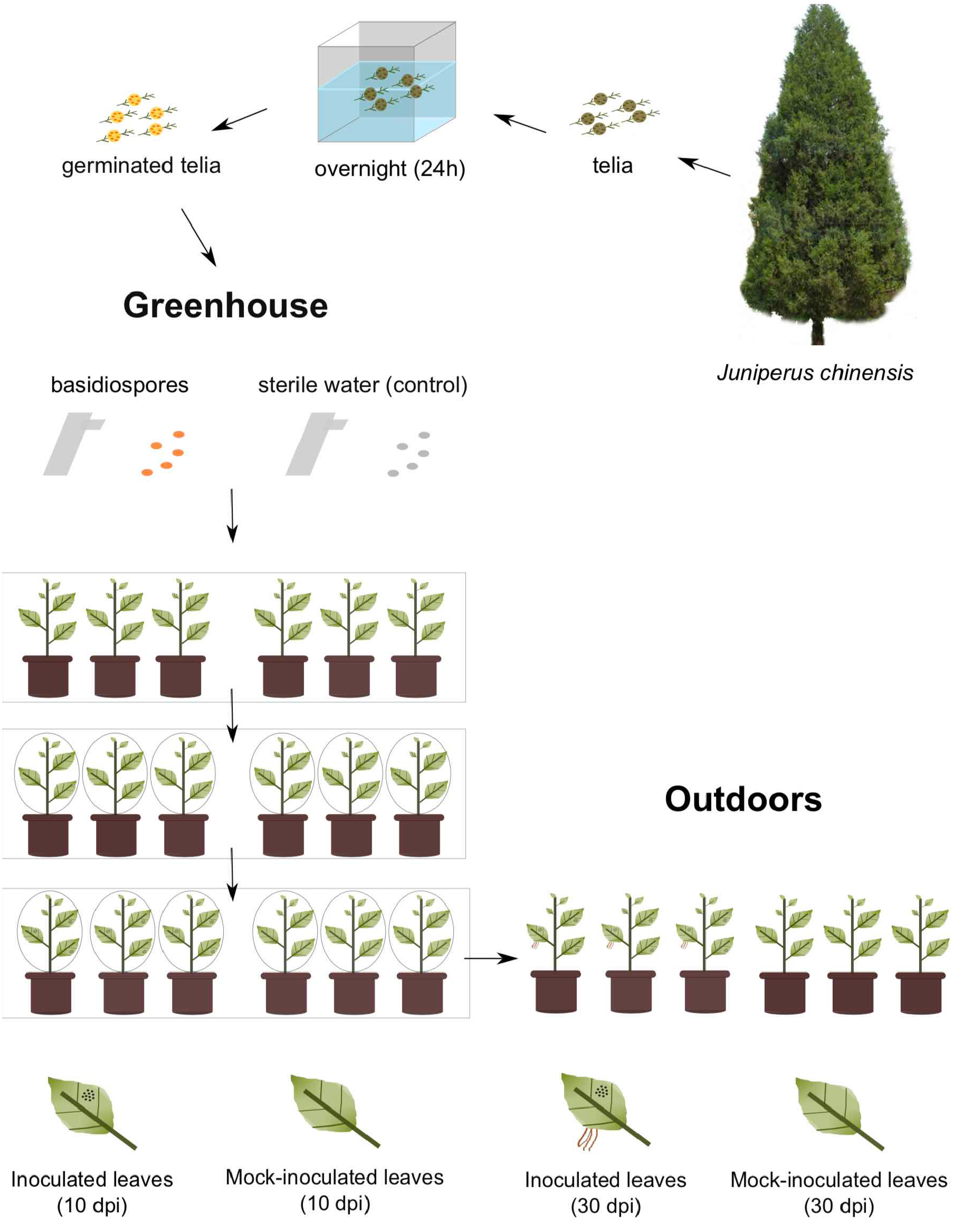
Experimental set-up established for artificial inoculation of apple seedlings by *Gymnosporangium yamadae*. Mature telia of *G. yamadae* were collected on wild *Juniperus chinensis* trees outdoor and were placed in water overnight to ensure germination and production of basidiospores. Two groups of 15 two-years-old apple seedlings were defined. The leaves of one group of seedlings were spray-inoculated with basidiospores and leaves of the other group were sprayed with sterile water as a mock-inoculated control. All the seedlings were covered with transparent plastic bags with openings at the top to ensure air exchange and were placed in a green-house with controlled temperature and moisture. Spermogonia formed on the upper surface of apple leaves after 10 days post inoculation (dpi). The infected and control seedlings groups were unbagged and moved outdoors. After 30 dpi, aecia formed on the lower surface of apple leaves. Samples were collected from leaves at 10 and 30 dpi. Random leave pieces of similar surface than in infected conditions were collected for the mock-inoculated controls at both time points.

**Figure S2.**
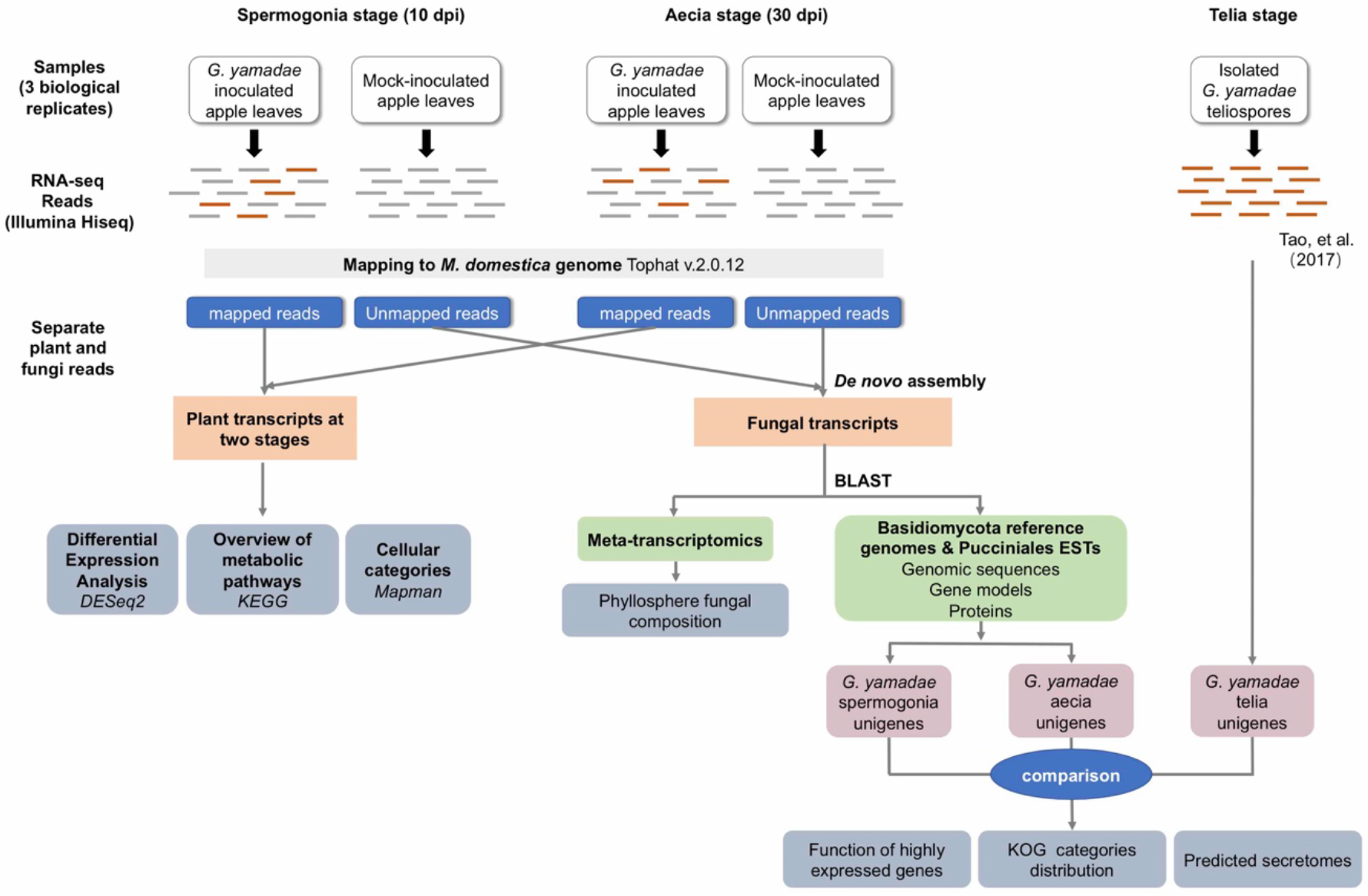
Overall RNA-seq and bioinformatic analysis pipeline used in the study. Briefly, after appropriate cleaning, reads obtained from Illumina RNAseq of three replicates in four conditions (rust-inoculated apple leaves and mock-inoculated controls at 10 and 30 days post-inoculation, dpi) were mapped to the apple reference genome to identify plant reads. Reads were assembled into unigenes at each of the four conditions and plant transcripts were compared to reference databases and pathways and between the inoculated and mock-inoculated conditions. Reads unmapped onto the apple genome were compared to reference fungal genomic databases to retrieve candidate apple rust fungus unigenes in the inoculated 10 and 30 dpi conditions before further annotation with ad hoc dedicated tools and databases. The fungal data were also compared to a previous dataset obtained at another apple rust fungal stage (telia; Tao et al. 2017). In parallel, non-plant reads were compared to global fungal databases through a dedicated meta-transcriptomic pipeline (see methods) in order to precisely assign transcripts to given fungal taxonomical levels.

**Figure S3.**
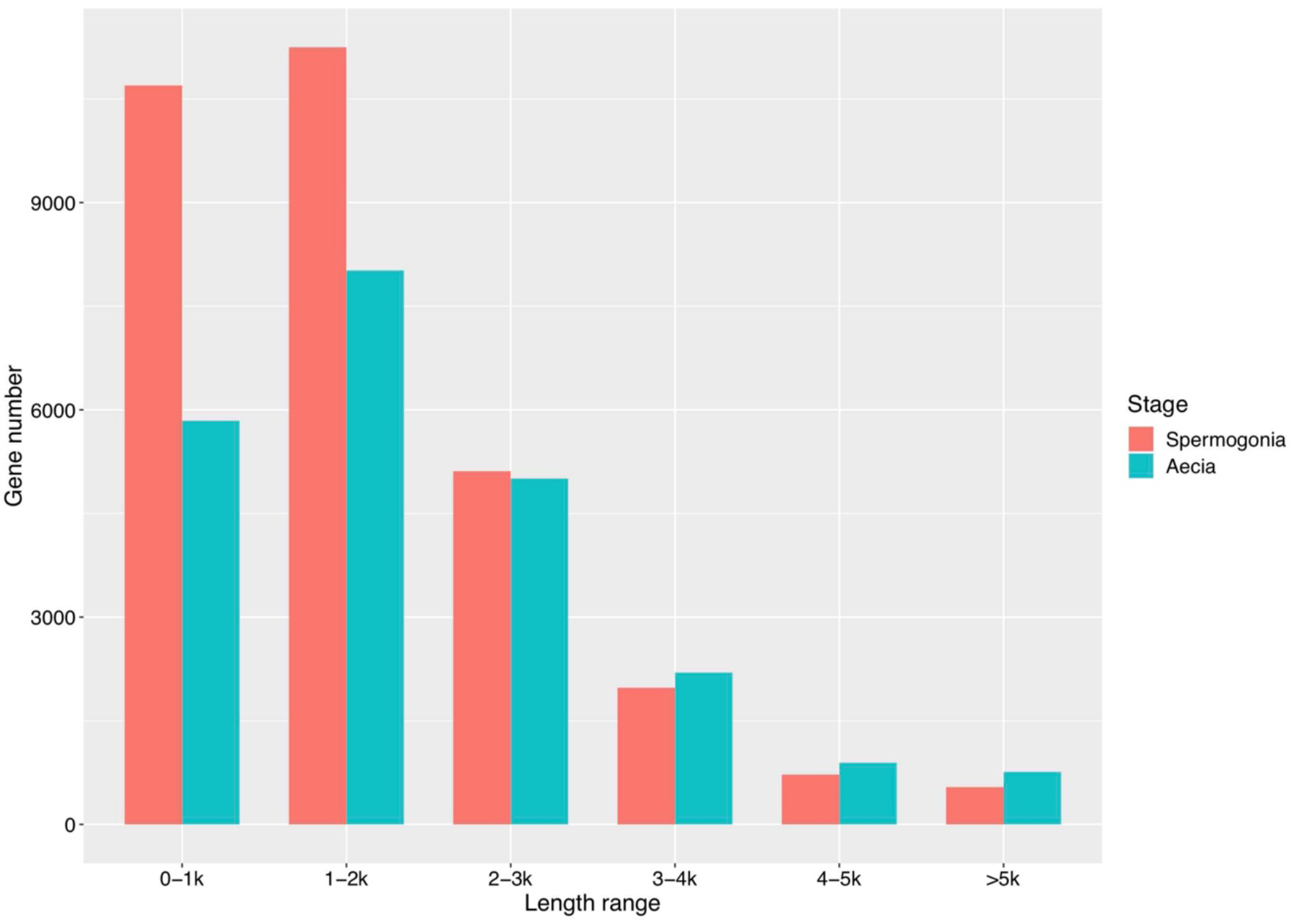
Length distribution of unigenes from spermogonia and aeica of *G. yamadae*.

**Figure S4.**
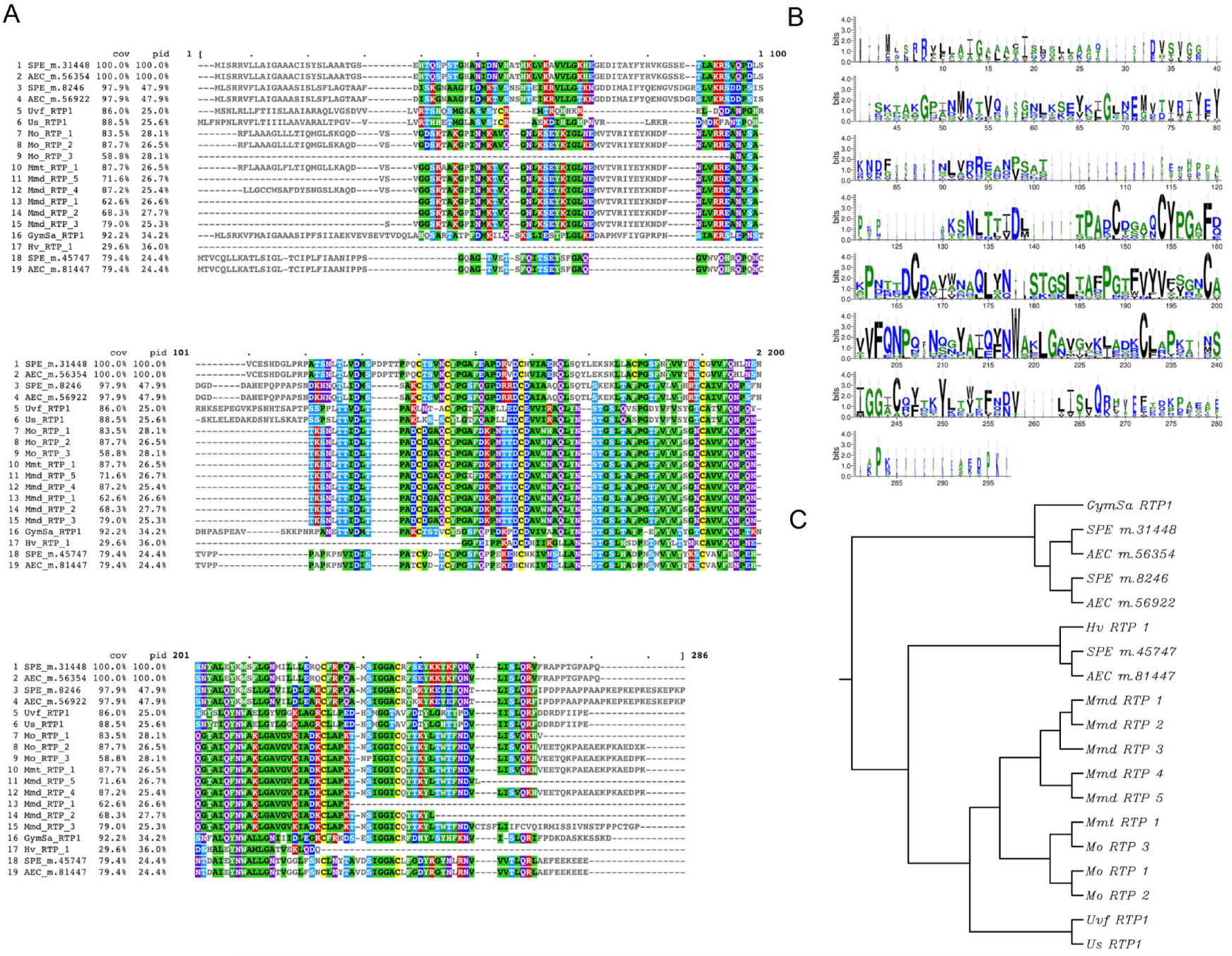
Multiple rust transferred protein 1 (RTP1) homologues from selected rust fungi. A: *Clustal Omega* alignment of rust proteins showing homology with RTP1 from *Uromyces fabae*. B: Conserved profile of selected proteins (Weblogo). C: UPGMA tree of RTP homologues sequences. Mo, *Melampsora occidentalis*; Mmt, *Melampsora medusae* f.sp. *tremuloidae*; Mmd, *Melampsora medusae* f.sp. *deltoidae*; Uvf, *Uromyces viviae-fabae*; Us, *Uromyces striatus*; Hv *Hemileia vastatrix*; GymSa, *Gymnosporangium sabinae*; SPE, *G. yamadae* spermogonia; AEC, *G. yamadae* aecia.

**Figure S5.**
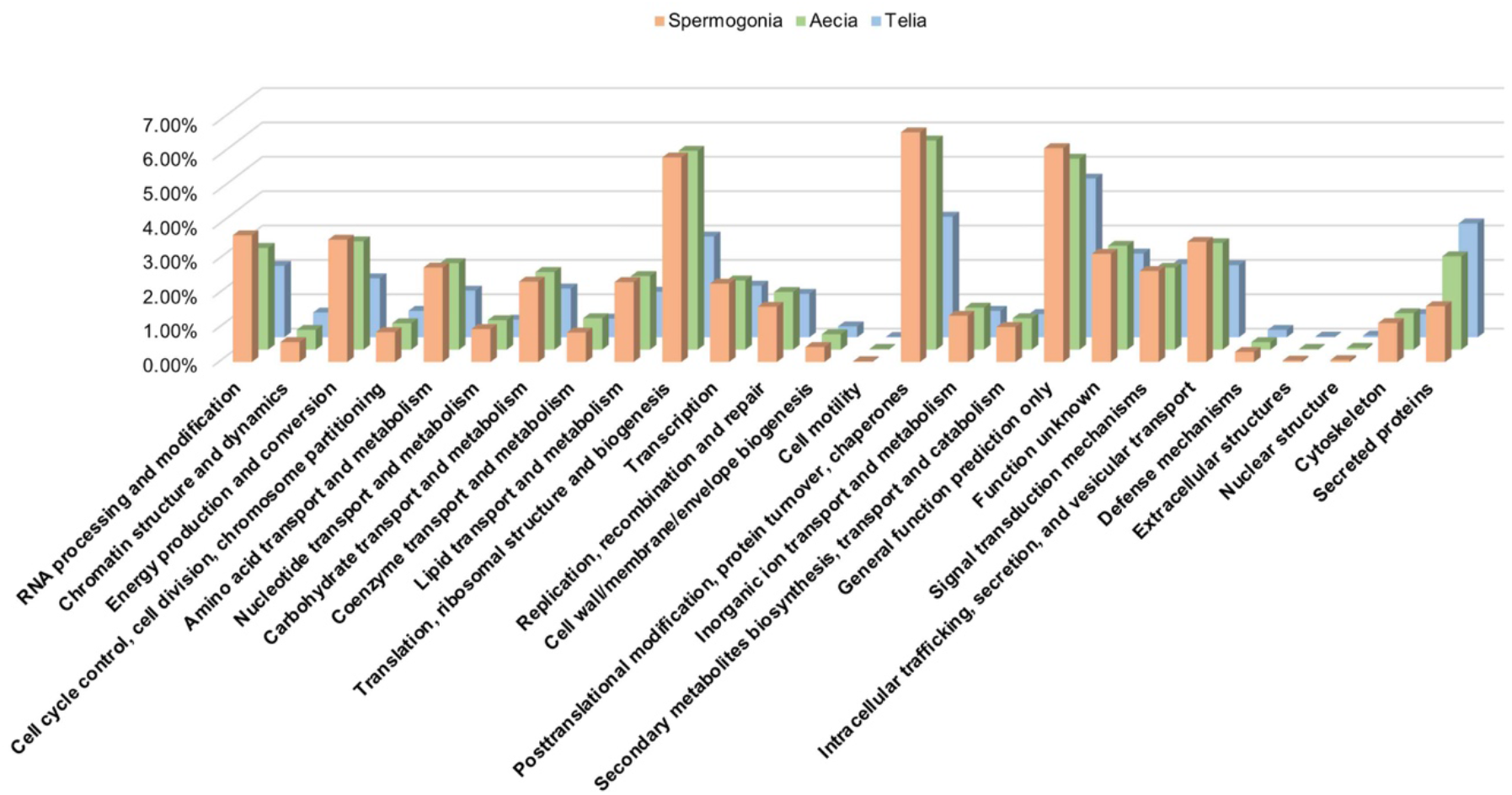
Distribution of unigenes from spermogonia, aecia and telia of *G. yamadae* annotated in eukaryotic orthologous group (KOG) categories. The unknown function category was removed for matters of figure readability, and the specifically added category “secreted proteins” refers to predicted secreted proteins of unknown function.

**Figure S6.**
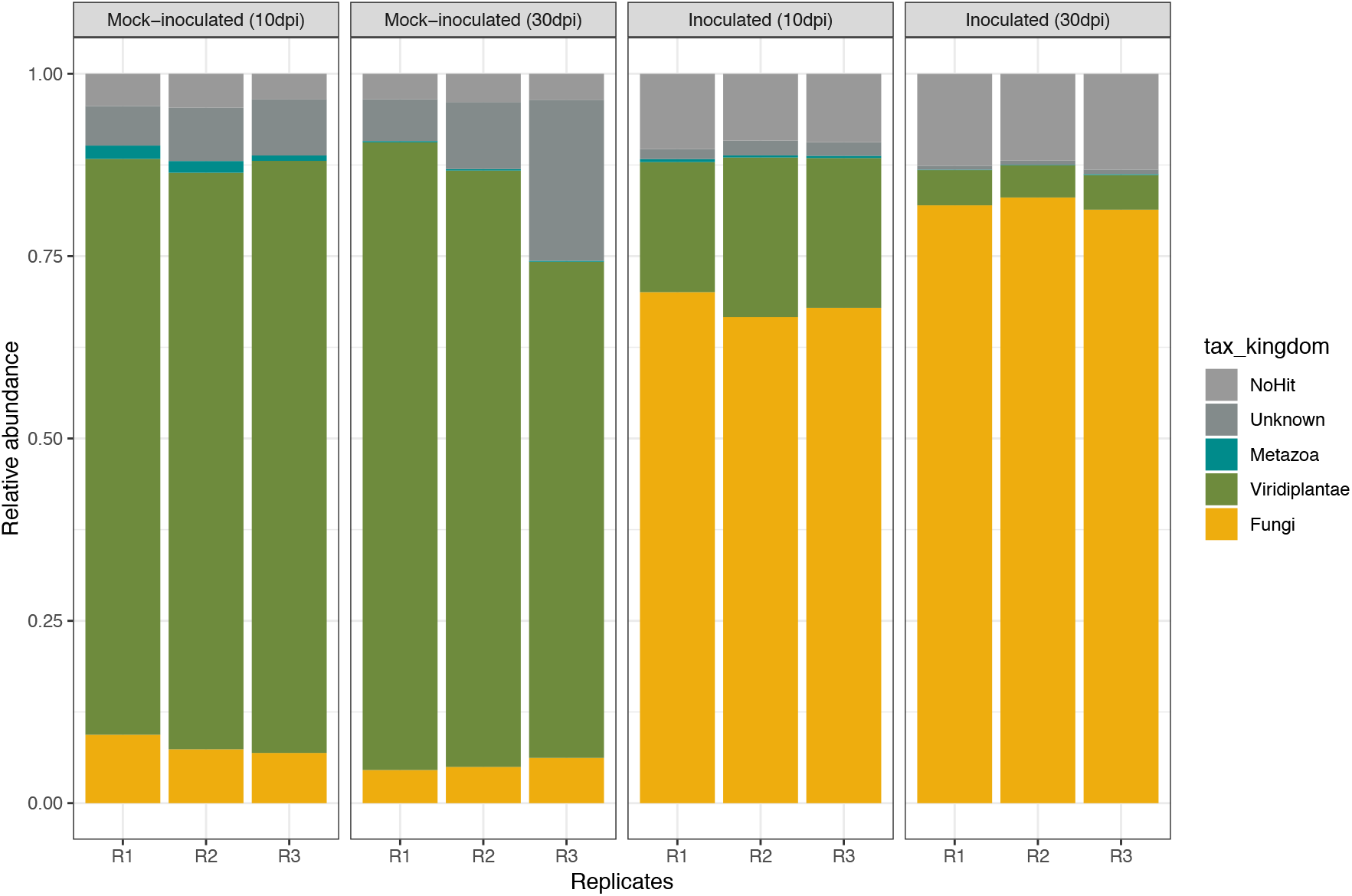
Apple phyllosphere community composition in rust inoculated and mock-inoculated conditions at 10 and 30 days post-inoculation (dpi). RNA-seq reads from apple leaves unmapped to the apple genome from each biological replicate (R1 to R3) were compared to a unigenes set built from the 12 replicates altogether. Non-fungal unigenes were annotated at the kingdom level by comparison to the NCBI non-redundant database. The figure presents the relative abundance of assigned reads in each replicate.

**Table S1.** *Malus domestica* transcripts expression in *Gymnosporangium yamadae* inoculated conditions and in mock-inoculated controls at 10 and 30 days post-inoculation (dpi) and assignment to metabolic pathways with subsequent enrichment analysis (MapMan and KEGG enrichment). Apple transcripts informations such as Gene_id, chromosome, Strand, Start, End, Length are provided. Read count and FPKM value for each gene were obtained for three biological replicates in four conditions (Inoculated_10dpi, Mock-inoculated_10dpi, Inoculated_30dpi, Mock-inoculated_30dpi) and differentially expressed levels between inoculated and mock-inoculated groups at 10 dpi and 30 dpi were evaluated with DESeq2 (adjusted *p*-value padj <0.05). All significantly differentially expressed genes between inoculated and mock-inoculated groups were assigned into MapMan functional categories and KEGG pathways. KEGG pathways enrichment tests realised with KOBAS v2.0 (hypergeometric test of KEGG pathways with Benjamini and Hochberg corrected *p*-values) are also presented.

**Table S2.** *Gymnosporangium yamadae* unigenes, predicted secreted proteins and identification of specific proteins at apple infection stages. For spermogonia and aecia unigenes, Gene id, Gene length, expression information (read count and FPKM value) for each biological replicate and annotation results from seven public databases (NR, NT, KO, Swissprot, PFAM, GO, KOG) are presented. For the secreted proteins predicted in spermogonia and aecia, CPL_fam represent CAZymes, proteases and lipases families predicted in the secretome; per C is the percentage of cysteine content in each amino acid sequence. NLS means Nuclear Localisation Signal prediction. Proteins from spermogonia, aecia and telia were clustered by Markov Cluster Algorithm (MCL) to identify the proteins specific to spermogonia, aecia and apple infection (spermogonia and aecia) stages.

**Table S3.** Meta-transcriptomic analysis results. Illumina RNA-seq trimmed reads (Total reads) and the un-mapped reads after mapping reads against the *Malus domestica* reference genome using CLC Genomics Workbench 11.0. All un-mapped reads from the 12 replicates in the four conditions were assembled together following a *de novo* co-assembly (Assembled unigenes) approach. The unigenes showing blast hits with JGI-Mycocosm fungal genomes are indicated as Fungal unigenes. The phyllosphere fungal composition for each replicate was analysed at the genus level by determining the relative abundance of reads mapped to the annotated fungal unigenes.

